# *Cryptosporidium* Oocyst Wall Proteins are true oocyst wall proteins, with COWP8 functioning to hold the inner and outer layers of the oocyst wall together

**DOI:** 10.1101/2024.07.20.604394

**Authors:** Ross Bacchetti, Sarah Stevens, Leandro Lemgruber, Mariana Azevedo Gonzalez Oliva, Emma M Sands, Konstantinos Alexandrou, Michele Tinti, Lee Robinson, Jack C Hanna, Simona Seizova, Peyton Goddard, Massimo Vassalli, Mattie C Pawlowic

## Abstract

Cryptosporidiosis is a significant cause of diarrhoeal disease contributing to substantial morbidity and mortality for the immunocompromised and for young children, especially those who are malnourished. There are no vaccines available and no effective treatments for these patients. Another challenge is that *Cryptosporidia* are waterborne and resistant to common water treatments including chlorination. *Cryptosporidia* are transmitted as an oocyst that is made up of a hardy oocyst wall that protects four parasites. Little is understood about how the oocyst is constructed, its composition, and the how it resists chlorination. A family of predicted *Cryptosporidium* Oocyst Wall Proteins (COWPs) was identified from the genome. Using a genetic approach, we confirm that all members of the COWP family localise to the oocyst wall. Our studies indicate that COWP2, 3 and 4 localise specifically to the oocyst “suture”, a zipper-like structure on the oocyst wall from which parasites emerge during infection. In parasites lacking COWP8, we observe that the inner and outer layers of the oocyst wall are no longer associated suggesting a role for COWP8 in oocyst wall morphology. Despite loss of COWP8, these transgenic parasites are viable, unchanged in mechanical strength, and retain resistance to chlorination. This work sets the foundation for future exploration of *Cryptosporidium* transmission.

## Introduction

Diarrhoeal disease is a major cause of morbidity and mortality for children under five years old, especially in low- and middle-income countries. The Global Enteric Multicentre Study revealed that cryptosporidiosis is the second leading cause of moderate to severe diarrhoeal disease.[1] Cryptosporidiosis was estimated to cause 8.2 million disability adjusted life years (DALYs) and 133,422 deaths in 2019.[2] Recurrent diarrhoeal episodes associated with cryptosporidiosis leads to alteration of gut morphology, growth stunting, delayed development, and impaired cognitive function.[3] There are no vaccines available, and the only drug approved to treat cryptosporidiosis is not effective in the populations that need it most—malnourished children and immunocompromised adults.[4]

Another major challenge of combatting this disease is the difficulty in removing it from the environment. *Cryptosporidium* is transmitted as an oocyst, an exceptionally hardy structure, granting protection to the four internal sporozoites. *Cryptosporidium* is a faecal-oral pathogen, and infection occurs by ingestion of contaminated food or water. *Cryptosporidia* are resistant to most water treatments including chlorination.[5] The ability to resist chemical disinfection contributes to cryptosporidiosis as the leading cause of reported waterborne outbreaks in high income countries[6], which are currently on the rise in the UK.[7] Outbreaks are common in recreational facilities that depend on chlorination for water treatment, daycare centres that care for young children, and in communities after large water treatment facilities fail, often due to increased influx of rainwater. Some species of *Cryptosporidium* also infect cattle and wildlife and can cause zoonotic infections.[8] Contribution of environmental contamination from animals to the contamination of water supplies for human consumption is unclear.

*Cryptosporidium* oocysts are slightly ovoid in shape and are approximately five microns in diameter.[9] The oocyst wall is made of three components: the outer oocyst wall, the inner oocyst wall, and the suture. The outer wall is SDS- and protease-resistant, made of acid-fast lipids, similar to the outer membrane of mycobacteria, and is sensitive to organic solvent treatment.[10, 11] This acid-fast lipid outer layer is hypothesised to act as a waxy coating that aids environmental survival, particularly protection from disinfectants and chlorination. The identity of the acid-fast lipids is still to be determined. The inner layer of the *Cryptosporidium* oocyst is composed of fibrillar glycoproteins. Once the oocyst wall is opened, the inner layer is susceptible to degradation by proteases.[10] The protein constituents of the inner wall are poorly characterised but appear to be highly crosslinked.[11, 12] It is hypothesised that these proteins offer structural strength and rigidity to the oocyst. The oocyst suture is a predefined opening, which spans approximately one-third to one-half the oocyst’s circumference, acting as the exit point for sporozoites.[13] Electron micrographs of the suture in cross-section reveal that the structure is formed from “zipper-like” teeth on the inside of the oocyst wall.[14] After the oocyst has been ingested, the suture opens allowing sporozoites to escape and invade the gut.[15, 16] The triggers for oocyst excystation and suture opening are unknown. However, the presence of enzymes and bile salts in the intestinal lumen, changes in CO_2_ concentration, changes in temperature, or the onset of reducing conditions have been implicated.[17]

There are few experimentally validated molecular markers for the outer and inner oocyst walls and there are no markers for the suture. The *Cryptosporidium* genome contains a family of nine predicted *Cryptosporidium* oocyst wall proteins, or COWPs.[18, 19] All COWPs have a signal peptide sequence, lack transmembrane domains, and are rich in cysteines. These cysteines occur at a high frequency whereby every 10th-12th amino acid is a cysteine residue. The abundance of cysteines in the COWPs suggest the possibility of forming inter- and intramolecular disulphide bonds, providing stability to the oocyst wall.[20]. COWP1 is experimentally confirmed to be expressed by macrogamonts in wall forming body organelles (WFB) and then secreted to form the oocyst wall.[21, 22] Antibodies generated against COWP8 suggest that this protein is also an integral inner wall protein.[19] The remaining COWPs have remained hypothetical oocyst wall proteins and their function in parasite transmission unexplored. Here, we demonstrate that all members of the COWP family are true oocyst wall proteins. This includes three proteins that localise to the suture, providing markers for this important structure. We investigated the function of the most abundant oocyst wall protein, COWP8, in oocyst wall structure and parasite transmission.

## Results

### COWPs localise to the oocyst wall and suture

To determine if COWP2-9 are oocyst wall proteins, CRISPR/Cas9 was used to generate reporter strains. Strains were engineered to have both a fluorescent protein and epitope tag, either mNeon-3×HA or mScarlet-I-3xmyc (**Figure 1A**). Since all COWPs contain a N-terminal signal peptide sequence, strains were designed as C-terminal fusions. Reporter strains were successfully generated for all COWPs (**Supplemental Figures 1-9**). An alternative tagging strategy was employed for COWP3-5 where a second copy of these COWPs was introduced at a non-essential locus (**Figure 1B**).[23] The respective promoters (entire upstream intergenic region) and open reading frame for COWP3-5 were cloned into the tagging constructs (mNeon-3xHA). CRISPR was used to insert these expression cassettes at *CpIMPDH*.

Oocysts were purified from faecal samples (**Figure 1C**) and examined by live fluorescence microscopy. COWP5-9 localise across the oocyst wall (**Figure 1 D-H**) in a similar pattern as previously observed for COWP1.[22] Additionally, immunoelectron microscopy confirmed localisation of COWP6 and COWP8 to the oocyst wall (**Supplemental Figure 5D and 7C**). Fluorescence microscopy indicates that COWP8 oocysts contain a few very small puncta in the space between sporozoites and the oocyst wall that are not associated with the residual body (**Figure 1E**). These were also observed by immunoelectron microscopy as globules of wall material secreted by the zygote that did not incorporate into the inner oocyst wall (**Supplemental Figure 7D**). Fluorescent signals for COWP7 and COWP9 were observed to be of lower intensity relative to COWP6 and COWP8 (**Figure 1** legend for image collection parameters). These data suggest COWP7 and 9 are expressed at a lower level that COWP1, 6 and 8 and is consistent with published transcriptomics data indicating that COWP1, COWP6, and COWP8 are the most abundant members of this family (**Supplemental Figure 10**).[22, 24]

Notably, COWP2-4 localise to the oocyst wall, but more specifically their localisation pattern is consistent with the suture, in both shape and length. COWP4 is located at the suture (**Figure 1J**). Unlike the other COWPs that localise evenly across the oocyst wall (COWPs 1, 5-9), COWP3 localises across the oocyst wall but is highly concentrated at the suture (**Figure 1K**). COWP2-mNeon localises to the suture, but also to a focus corresponding to the residual body (**Figure 1L**). We hypothesise that the small puncta observed in COWP2 and COWP4 are similar to the structures observed in the COWP8 strain. The relatively low protein abundance of COWP2 and COWP4 prevented observation of labelling by immunoelectron microscopy.

Application of genetics and microscopy confirms that all members of the COWP family are true oocyst wall proteins. Further, we discovered that COWP2-4 specifically localise to the *Cryptosporidium* oocyst suture, providing the first markers for this structure.

**Figure 1.**
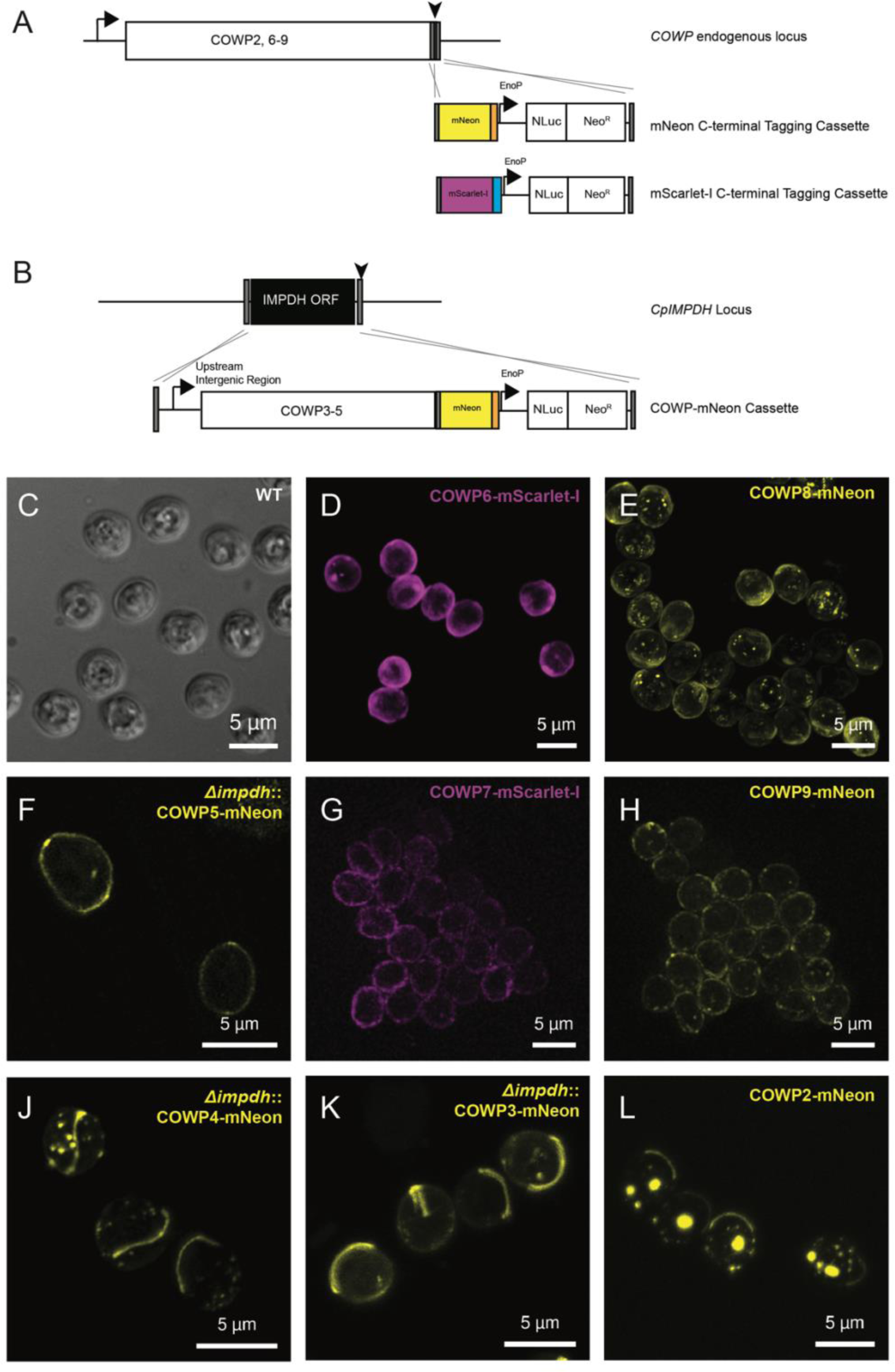
*Cryptosporidium* Oocyst Wall Protein (COWP) family are confirmed oocyst wall proteins. **A**) CRISPR/Cas9 was used to produce strains of *Cryptosporidium* where COWP2, 6-9 are individually fused at their C-terminus to a fluorescent protein, either mNeon (yellow) or mScarlet-I (magenta). **B**) Expression of COWP3-5 tagged at the C-terminus with mNeon were driven by their native promoter and targeted for integration at the *CpIMPDH* locus. **C**) Live microscopy of wild type *Cryptosporidium parvum* oocysts. **D-L**) Fluorescence microscopy of live oocysts confirms that COWPs localise to the oocyst wall and/or suture. Images collected, as indicated, using a Deltavision widefield epifluorescence microscope, or on a Zeiss LSM880 Airyscan microscope either in confocal or super resolution mode (Airyscan mode). Representative images shown; contrast adjusted individually for each image and not normalised. Image collection parameters for each image: **C**) widefield epifluorescence microscope, single z-plane, exposure time 0.08 msec, laser 50%. **D**) confocal, projection of 29 z-planes, exposure time 0.01 msec, laser power 2%. **E**) super resolution, projection of 20 z-planes, exposure time 0.038 msec, laser power 2.2%. **F**) super resolution, single z-plane, exposure time 0.043 msec, laser power 2.2%. **G**) widefield epifluorescence microscope, single z-plane, exposure time 0.5 sec, laser 50%. **H**) widefield epifluorescence microscope, projection of 12 z-planes, exposure time 0.5 sec, laser 50%. **J**) super resolution, 39 z-planes, exposure time 0.043 msec, laser power 2.3%. **K**) confocal, projection of 10 z-planes, exposure time 0.02 msec, laser power 2%. **L**) confocal, projection of 16 z-planes, exposure time 0.02 msec, laser power 2%.

### COWP6 and COWP8 are expressed by macrogamonts and are secreted after fertilisation to form the oocyst wall

To determine which parasite life cycle stages express COWPs, reporter strains were cultured *in vitro* or *in vivo* and expression of COWPs in different life cycle stages was observed using fluorescence microscopy. We focussed on COWP6 and COWP8 as the expression was high enough to confidently observe by fluorescence microscopy; COWP7 and COWP9 were examined, but expression could not be visualised by fluorescence microscopy, likely due to low expression levels.

We lack markers for discrete life cycle stages and use shape and number of nuclei to differentiate life cycle stages. Expression of COWP6 and COWP8 was undetectable in asexual stages including single nuclei trophozoites (**Supplemental Figure 11A**) and eight-nuclei meronts (**Supplemental Figure 11B**). Males (microgamonts), which have up to 16 bullet-shaped nuclei, also do not express COWP6 or COWP8 (**Supplemental Figure 11C**).

COWP6 and COWP8 expression was exclusively observed in females, called macrogamonts (**Figure 2A and C**). Macrogamonts have a single nuclei and synthesise the components of the oocyst wall as they mature.[22] These components are stored in organelles called “wall forming bodies” (WFB) which have a punctate pattern.[21] Previously, COWP1 was localised to wall forming bodies by immunoelectron microscopy.[21] Recent examination of COWP1-mNeon parasites identified a similar punctate localisation consistent in size, shape, and number with WFBs.[22]

We observe the same WFB-like localization pattern of our COWP6 and COWP8 reporter strains. Prior to fertilisation (characterised by a single large nuclei), macrogamonts parasites express COWP6 (**Figure 2A**) and COWP8 (**Figure 2C**) in numerous puncta of a similar, and small size. After fertilisation and meiosis (four nuclei) COWPs are secreted to form the oocyst wall (**Figure 2B and 2D**).

**Figure 2.**
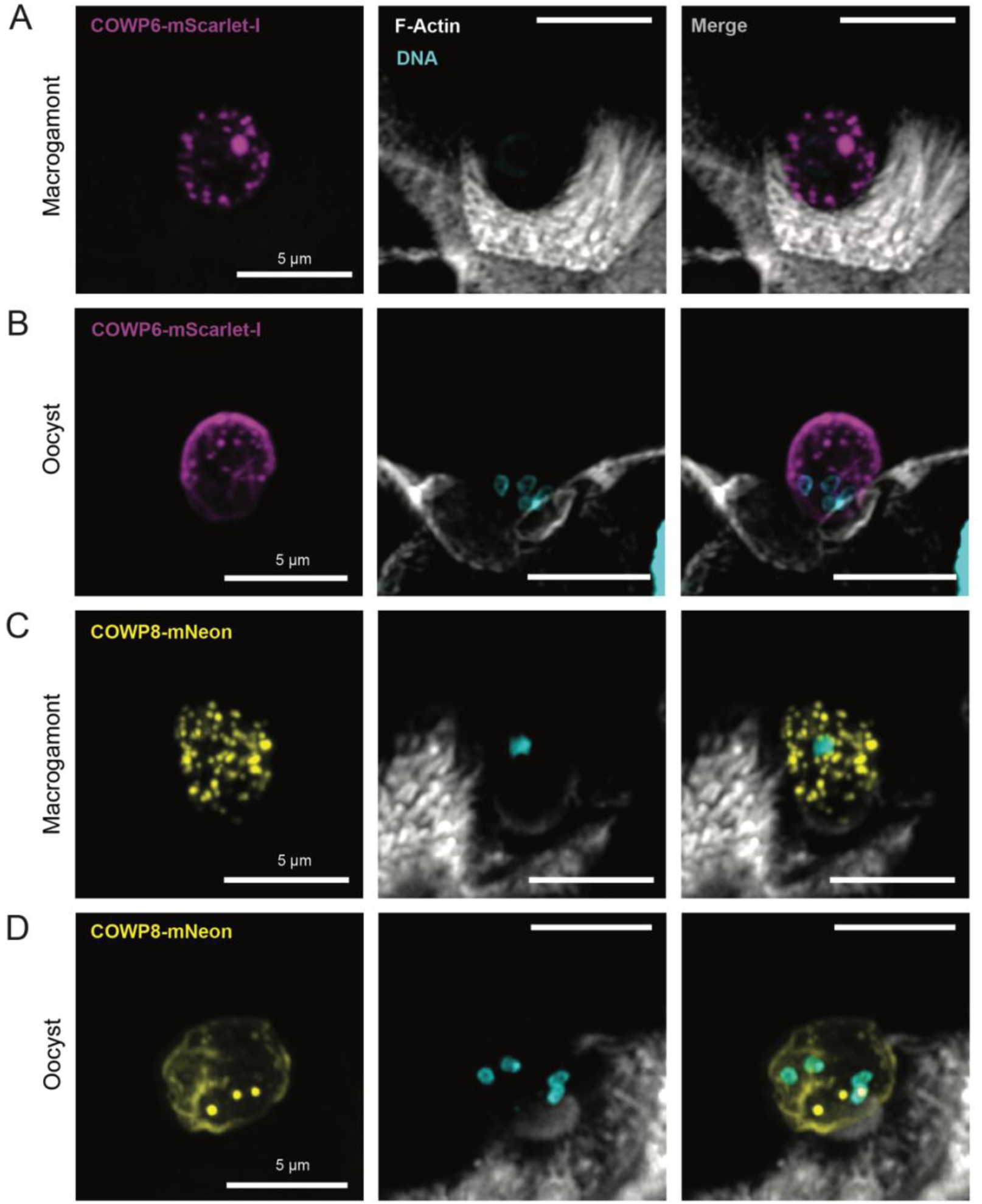
COWPs are expressed by macrogamonts and are secreted to form the oocyst wall. Super resolution microscopy of ileal tissue collected from IFN-γ knockout mouse infected with COWP6-mScarlet (**A** and **B**) or COWP8-mNeon (**C** and **D**) strains. Mice were culled at peak infection and tissue was processed for histology. Macrogamonts are identified by size (∼ 5 µm) and a single nucleus; oocysts are identified similarly by size and presence of 4 nuclei. Brush border (F-actin stained with phalloidin-AlexaFluor 647, white) and nuclei (DNA stained with Hoechst, cyan). **A**) Prior to fertilisation, COWP6 (magenta) is expressed in numerous puncta by the female parasites, resembling wall forming body organelles. **B**) After fertilization, COWP6 is secreted to form the oocyst wall. **C**) COWP8 (yellow) is expressed in puncta similar in size and number to COWP6, and **D**) after fertilisation is secreted to form the oocyst wall. Small puncta of COWP8-mNeon remain after oocyst wall formation, consistent with live microscopy of oocyst purified from faecal material (reported in Figure 1D). Images collected on a Zeiss LSM880 Airyscan microscope, Airyscan mode. Representative images shown; contrast adjusted individually for each image and not normalised. Scale bar for all images is 5 µm. Image collection parameters for each image: **A**) projection of 5 z-planes, exposure time mScarlet-I 0.028 msec and laser 2%, exposure time Alexa-647 0.024 msec and laser 0.5%, exposure time Hoescht 0.010 msec and laser 3%. **B**) projection of 10 z-planes, exposure time mScarlet-I 0.028 msec and laser 2%, exposure time Alexa-647 0.024 msec and laser 0.5%, exposure time Hoescht 0.010 msec and laser 3%.**C**) projection of 9 z-planes, exposure time mNeon 0.023 msec and laser 2.4%, exposure time Alexa-647 0.020 msec and laser 0.5%, exposure time Hoescht 0.022 msec and laser 3%. **D**) projection of 17 z-planes, exposure time mNeon 0.020 msec and laser 2.4%, exposure time Alexa-647 0.020 msec and laser 0.5%, exposure time Hoescht 0.022 msec and laser 3%.

### COWP8 is not essential for parasite survival or transmission

To understand the role of COWPs in construction of the oocyst wall and parasite transmission, we used CRISPR/Cas9 to generate parasites lacking COWP8. We focussed on COWP8 as it is the most highly expressed member of the COWP family. From transcriptomics data, COWP8 is in the 99% percentile of genes expressed by macrogamonts and is expressed 1.5x as the next most abundant COWP, COWP6 (**Supplemental Figure 10)**.[22]

The repair cassette was designed to replace COWP8 with an mScarlet-I fluorescent protein and the NLuc-Neo^R^ fusion protein, driven by a strong, constitutive promoter (**Supplemental Figure 12**). *Δcowp8* parasites were viable *in vivo* (**Figure 3A-F**) and their infection pattern followed a typical acute pattern, indistinguishable from the pattern observed for passage of COWP8-mNeon reporter strain (**Figure 3A-F**). Infection was unaffected by type of inoculum, as both a faecal sample (**Figure 3B**) or purified oocysts (**Figure 3C**) generated robust infection. There was no observable difference between the morphology of wild type (**Figure 3G and H**) and *Δcowp8* oocysts by confocal microscopy (**Figure 3J and K**). Parasite burden of *Δcowp8* infected tissue was observed using fluorescent microscopy. Intestinal tissue was collected from mice at peak infection levels (determined by faecal NLuc) and imaged. High levels of infection were observed (**Figure 3L**) along the length of the villi. We conclude that COWP8 is not essential for parasite survival, and loss of this protein does not affect infectivity or impair *in vivo* transmission under standard laboratory conditions.

**Figure 3.**
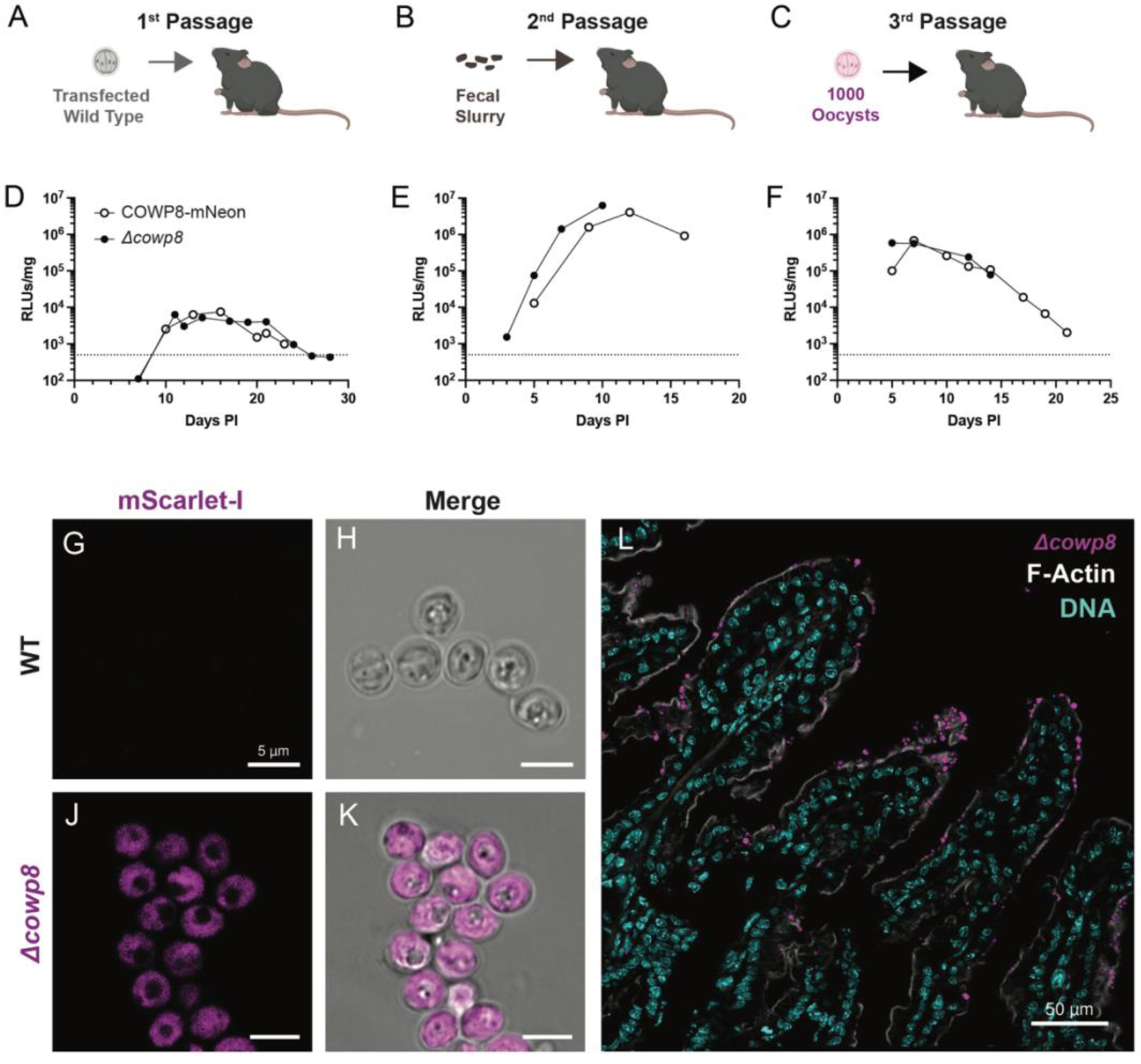
COWP8 is not essential for parasite growth and transmission. Experimental design for passage of *Δcowp8* strain in interferon-gamma knockout mice. **A**) Initial passage where inoculum is wild type sporozoites transfected with CRISPR machinery to target COWP8 for deletion (**Supplemental Figure 12A**), **B**) second passage infected with slurry made of faecal material from first passage, **C**) and third passage infected with oocysts purified from the second passage. Infection level of COWP8-mNeon (white circles) and *Δcowp8* strains (black circles) is similar and follows typical acute infection pattern in **D**) first, **E**) second, and **F**) third passages. Infection level of mice as determined by faecal NLuc, limit of detection at 500 RLU/mg, dotted line. Average and SD of three technical replicates of one biological replicate. Confocal microscopy of live wild type (**G** and **H**) and *Δcowp8* (**J** and **K**) confirms mScarlet-I cytoplasmic expression of *Δcowp8* strain (magenta); scale bar 5 µm. **L**) Super resolution microscopy of tissue collected from interferon-gamma knockout mouse infected with *Δcowp8* culled at peak infection and processed for histology. Brush border (F-actin stained with phalloidin, white) of ileal villi (DNA stained with Hoechst, cyan) are highly infected with *Δcowp8* (mScarlet-I expressed in the parasite cytoplasm, magenta). Images collected on a Zeiss LSM880 Airyscan microscope, (Airyscan mode for **L**). Representative images shown.

### COWP8 functions to hold the inner and outer wall together

We visualised changes to the oocyst wall morphology by scanning electron microscopy (SEM), and changes to the inner and outer layers of the wall using transmission electron microscopy (TEM). By SEM, COWP8-mNeon and *Δcowp8* cannot be distinguished. This is consistent with confocal microscopy which did not identify morphological changes. Further investigation by TEM revealed that loss of COWP8 causes the inner and outer layers of the oocyst wall to disassociate (**Figure 4A and B; Supplemental Figure 14 D and F**). There also appears to be more space between sporozoites and the inner wall. This was not observed by TEM for any COWP8-mNeon oocysts, which were prepared from faecal material and processed in parallel with *Δcowp8* oocysts. These observations suggest that COWP8 may function as a “glue” that holds the inner and outer layers of the oocyst wall together.

Oocysts were analysed by quantitative mass spectrometry to investigate if parasites can compensate for loss of COWP8 by upregulating other members of the COWP family. WT and *Δcowp8* oocysts were excysted, proteins were extracted, samples were labelled with tandem mass tags, and then analysed using quantitative mass spectrometry. The expression of COWPs are not significantly changed upon loss of COWP8. This is consistent with the observation that although the layers of the oocyst wall are no longer physically connected in *Δcowp8* oocysts, both inner and outer layers appear unchanged (**Supplemental Figure 12E**). A small number of differentially expressed proteins were observed in *Δcowp8* but are not specifically related to oocyst biology (**Supplementary** Figure 13**; Supplemental Table 4**).

### COWP8 mutants retain mechanical strength

To determine if there are changes to oocyst strength upon loss of COWP8, we measured *Δcowp8* oocyst walls using a biomechanical approach called NanoIndentation. This approach is commonly used to measure the hardness and elastic modulus of biological materials.[25] Oocysts are immobilised on a petri dish and are probed under a microscope. The probe approaches the sample at a constant speed, generating a force-displacement curve as the probe indents the sample with load and indentation depth. Young’s Modulus, which is the mechanical property measuring stiffness of a material by defining the relationship between stress and strain, can be obtained by fitting force-displacement curves to a corresponding mathematical model. Oocysts were immobilised on a dish and subjected to NanoIndentation using a Chiaro NanoIndentor. From raw force displacement curves (**Supplemental Figure 15A**), force indentation plots were created using average force measurements obtained from 0-100 nm of indentation (**Supplemental Figure 15B and C**). Young’s Modulus was calculated by fitting the force indentation plots to the Hertz mathematical model for both wild type and *Δcowp8* oocysts. No difference in mechanical strength was observed (**Figure 4C**) and wild type and *Δcowp8* oocysts were determined to have an average Young’s Modulus of 1.2 Megapascals (MPa).

Although COWP8 is not required for infection of immunocompromised mice under laboratory conditions, it could be required for survival in the environment. Oocysts are resistant to treatment with chlorination and can survive exposure to high levels of bleach. Purified COWP8-mNeon and *Δcowp8* oocysts were treated with 8% bleach, washed thoroughly, and used to infect mice. No difference in infection levels was observed for *Δcowp8* parasites (**Supplemental Figure 14A**), confirming that oocysts remain resistant to chlorination despite loss of COWP8. In addition to their resistance to disinfectants, *Cryptosporidium* oocysts can persist and survive for considerable lengths of time. Oocysts stored below 15 °C remain highly infectious even after six months.[26] Parasite infectivity begins to be greatly compromised only when oocysts are stored at temperatures above 37 °C.[5, 27] We tested the effect of heat treatment on oocysts lacking COWP8. Treatment at 60 °C for 5 minutes was sufficient to render both COWP8-mNeon and *Δcowp8* non-infectious (**Supplemental Figure 14B**).

**Figure 4.**
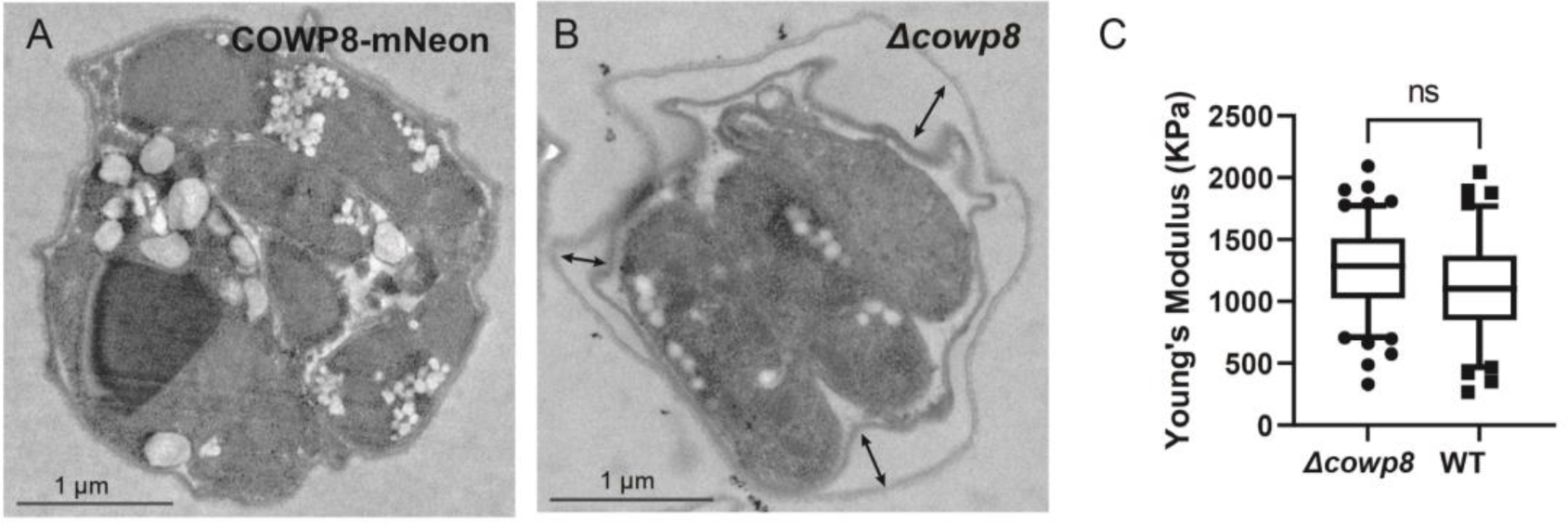
COWP8 links inner and outer walls but does not alter mechanical strength of the oocyst wall. Transmission electron microscopy was performed on **A**) COWP8-mNeon and **B**) *Δcowp8* oocysts revealing a “gap” between inner and outer oocysts walls (indicated by arrows) upon loss of COWP8. **C**) Wild type and *Δcowp8* oocysts were subjected to NanoIndentation to measure differences in the mechanical strength of the oocyst wall. Young’s modulus was obtained from fitting the revised indentation plot (**Supplemental Figure 15C**) to the Hertz mathematical model, producing box and whisker plots (10-90 percentile). Statistical analysis carried out on obtained data (Welch’s t-test) concluded that there is no significant difference in mechanical strength between wild type and *Δcowp8* oocysts (*p* >0.05).

## Discussion

Despite their name, few members of the *Cryptosporidium* Oocyst Wall Protein family were experimentally confirmed as oocyst wall proteins.[19] Using CRISPR, COWPs 2-9 were fluorescently tagged at the C-terminus and their localisation in the oocyst wall was validated by fluorescent microscopy. Although tagging was attempted several times for COWPs 3, 4, and 5 (multiple independent transfections using both florescent protein tag and smaller epitope tags), these strains were not recovered (**Supplemental Figure 9**). It is probable that introduction of the tagging construct at the endogenous locus disrupted expression of the neighbouring gene and was not tolerated. Expression of these tagged COWPs using their corresponding promoter at a secondary location in the genome was well tolerated.

Interestingly, COWPs 2, 3, and 4 localised to a structure consistent with the size and shape of the suture. The suture forms a seam along the shorter end of the slightly oval-shaped oocyst (**Figure 5A**). During the process of excystation, the suture opens allowing sporozoites to exit and infect the gut. COWP 2, 3, 4 are newly identified markers for this important structure and will allow further investigation of the process by which the suture is constructed and how it is “opened” during excystation.

For the more highly expressed family members, COWP6 and COWP8, their exclusive expression in macrogamonts was confirmed. Like COWP1, COWP6 and COWP8 are expressed in multiple puncta throughout the cytoplasm of the macrogamont. These structures resemble wall forming body organelles. New selection markers present on opportunity to create a reporter strain where more than one COWP is tagged, allowing colocalization studies.[28, 29]

COWP8 is the most highly expressed of the COWP family and is expressed last. This high and late level of expression suggest a unique role in oocyst construction and parasite transmission. Transcriptomics from single cell RNA sequencing data [24] reveal that COWP8 is the last of the COWPs to be expressed. There are six developmental clusters for macrogamonts. While all the other COWPs are expressed starting in the fourth cluster, COWP8 is exclusively expressed only in the final cluster. For that reason, we used CRISPR to attempt to knockout COWP8. Surprisingly, *Δcowp8* parasites are viable and have no transmission defect. Quantitative proteomics determined that *Δcowp8* parasites do not upregulate expression of other members of the COWP family to compensate for loss of COWP8.

Like wild type oocysts, *Δcowp8* are resistant to bleach treatment but are susceptible to heat treatment. Confocal microscopy did not reveal morphological changes to the oocyst walls of *Δcowp8* parasites. By transmission electron microscopy we observed that the inner and outer walls of *Δcowp8* are no longer attached. We failed to observe this phenotype for any COWP8-mNeon oocysts, which were prepared from faecal samples, embedded, sectioned, and examined by electron microscopy in parallel. Future work will quantify this phenotype. Although COWP8 is depleted, the inner and outer layers are maintained at thickness observed in wild type. Biomechanical measurements of *Δcowp8* determined the strength to be similar to wild type. Although the layers are no longer attached, they are still functional to protect the parasites in the environment (*Δcowp8* oocysts are viable for transmission, resistant to bleach, and physically strong).

We hypothesise that COWP8 likely serves as a “glue” holding together the outer layer of the oocyst wall, made of acid-fast lipids, and the inner protein-rich layer (**Figure 5B**). COWP8 may be expressed high and late in order to form this “glue” at the right time during oocyst wall construction. Future work will investigate the mechanisms by which COWP8 interacts with each layer.

**Figure 5.**
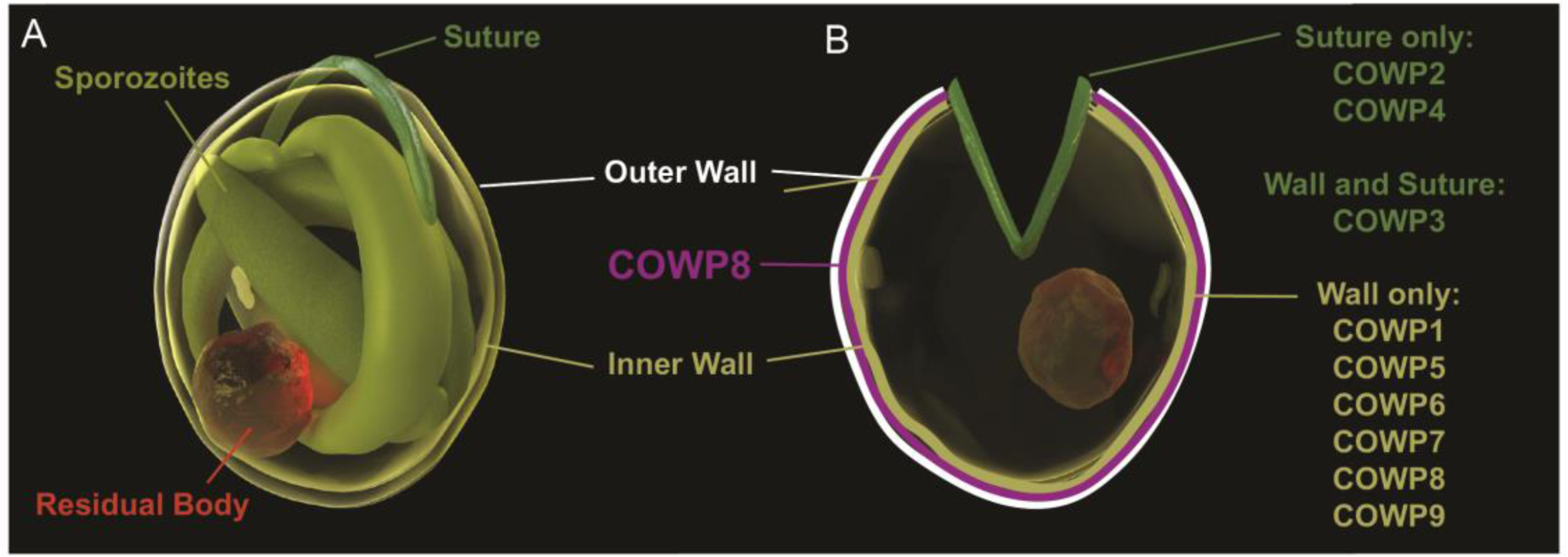
Model of COWPs in oocyst wall structure. **A**) 3D rendering of the *Cryptosporidium* oocyst structure, where 4 sporozoites (olive green) and a residual body (red) are tightly packed inside the slightly oval shell of the oocyst wall. Inner and outer walls are sealed closed by the suture (green). **B**) Upon excystation the suture opens, allowing sporozoites to escape often leaving behind the residual body inside the excysted oocyst wall. COWPs 1, 5, 6, 7, 8, 9 localise to the oocyst wall only. COWP2 and COWP3 localise to both the oocyst wall and the suture, while COWP4 is localised to the suture only. COWP8 (purple) functions to hold the inner (yellow) and outer (white) layers of the oocyst wall together. Illustrations by Konstantinos Alexandrou.

## Material and Methods

### Materials

Wild type *Cryptosporidium parvum* oocysts (Iowa strain) were purchased from Bunchgrass Farms and stored at 4°C in PBS (Deary, Idaho, USA). Oligonucleotides produced by either the University of Dundee Oligonucleotide Synthesis Service or Sigma-Aldrich (**Supplemental Table 1**). mScarlet-I described throughout is codon optimized for *C. parvum* (GeneScript; sequences available in **Supplemental Table 2**).

### Molecular Cloning and DNA Sequencing

PCR performed using high-fidelity DNA Polymerase (Primestar Max, Takara) and cloning with NEBuilder HiFi DNA Assembly (NEB). DNA Sequences confirmed by sequencing by Genewiz, University of Dundee MRC PPU Reagents and Services, or Plasmidsaurus. All plasmids created for this work listed in **Supplemental Table 2** with links available for DNA sequences.

Construct used for C-terminal tagging with mNeonGreen-3xHA previously described, referred to as mNeon throughout.[28] A second construct was created with mScarlet-I fused to a 3xc-myc epitope tag followed by an NLuc-Neo^R^ fusion under the control of constitutively expressed *CpEnolase* promoter to generate “pLIC-mScarlet-I-3xc-myc_aldo 3’ ’ utr_ENNE”. Construct for deletion of *CpCOWP8* contains mScarlet-I, 2A linker, and an NLuc-Neo^R^ fusion, all of which is under the control of the *CpEnolase* promoter to generate “EnoP_mScarlet-I-ENNE”.

Due to the location of the gRNA at the C-terminus, the last 129 bp of *CpCOWP4* was cloned into a tagging construct fused directly to the mNeon tagging construct. The codon for cysteine at amino acid position 845 was mutated from “TGC” to “TGT” to prevent further cutting by Cas9 to generate “*Cp*COWP4 C-term-mNeon-3xHA_ENNE”. Similarly, the last 114 bp of *CpCOWP9* was cloned into a tagging construct fused directly to an mNeon-3xHA. The codon for lysine at amino acid position 1180 was mutated from “AAG” to “AAA” to prevent further cutting by Cas9 to generate “*Cp*COWP9 C-term-mNeon-3xHA_ENNE”.

To express *CpCOWP3*, *CpCOWP4*, *CpCOWP5* under their endogenous promoter from an ectopic location, the entire intergenic region upstream of the corresponding COWP gene and its full open reading frame (except the stop codon) was amplified using PCR from *C. parvum* genomic DNA. This was cloned to generate “Cowp3P_COWP3ORF-mNeon-3xHA_ENNE,” “Cowp4P_COWP4ORF-mNeon-3xHA_ENNE,” and “Cowp5P_COWP5ORF-mNeon-3xHA_ENNE”.

Guide oligonucleotides were designed for *COWPs* 2-9 (**Supplemental Table 1**) and ligated into a plasmid containing Cas9/*Cp*U6 promoter containing plasmid [30] using restriction enzyme cloning as previously described [31]. To generate the *Dimpdh::COWP3-mNeon*, *Dimpdh::COW4-mNeon*, and *Dimpdh::COWP5-mNeon*, a guide for the inosine monophosphate dehydrogenase (IMPDH cgd6_20) locus was used [23].

### CRISPR Design

Linear repair cassettes were generated via PCR using PrimeSTAR MAX DNA polymerase (Takara). Fifty bp of homology, typically directly flanking the site targeted by CRISPR, was incorporated on primer.

To generate COWP2-mNeon, *CpCOWP2* (cgd7_1800) was targeted by a gRNA located 38 bps upstream of the stop codon. Regions of homology consisting of the 68 bp directly upstream of the stop codon, and 50 bp located 116 bp downstream of the stop codon. A synonymous mutation of serine at position 1373 encoding the PAM (AGG to AGA) was engineered to prevent further targeting of Cas9 upon integration of the repair cassette into the genome.

To attempt to generate COWP3-mNeon and COWP3-HA, *CpCOWP3* (cgd4_670) was targeted by a gRNA located 16 bps after the stop codon. Regions of homology consisting of the 50 bp directly upstream of the stop codon, and 50 bp located 177 bp downstream of the stop codon.

To attempt to generate COWP4-mNeon, *CpCOWP4* (cgd8_3350) was targeted by a gRNA located 124 bps after the stop codon. Regions of homology consisting of the 179 bp upstream of the stop codon, and 50 bp located 153 bp downstream of the stop codon.

To attempt to generate COWP5-mNeon and COWP5-HA, *CpCOWP5* (cgd7_5150) was targeted by a gRNA located 17 bps upstream of the stop codon. Regions of homology consisting of the 71 bp upstream of the stop codon, and 50 bp located 130 bp downstream of the stop codon. A synonymous mutation of threonine at position 623 encoding the PAM (CGG to CTG) was engineered to prevent further targeting of Cas9 upon integration of the repair cassette into the genome.

To generate *Δimpdh::COWP3-mNeon*, *Δimpdh::COWP4-mNeon*, and *Δimpdh::COWP5-mNeon* the *CpIMPDH* locus was targeted using CRISPR as previously described [23].

To generate COWP6-mScarlet-I, *CpCOWP6* (cgd4_3090) was targeted by a gRNA located 13 bps downstream of the stop codon. Regions of homology consisting of the 57 bp directly upstream of the stop codon, and 50 bp located 54 bp downstream of the stop codon. A synonymous mutation of serine at position 536 encoding the PAM (AGC to AGT) was engineered to prevent further targeting of Cas9 upon integration of the repair cassette into the genome.

To generate COWP7-mNeon, *CpCOWP7* (cgd4_500) was targeted with a guide RNA located 19 bp upstream of the stop codon. Homology regions of 92 bp located directly upstream from the stop codon, and 50 bp located 56 bp downstream of the stop codon. The PAM was mutated such that proline at position 827 was replaced with alanine.

To generate COWP8-mNeon, *CpCOWP8* (cgd6_200) was targeted with a guide RNA found 39 bp upstream of the stop codon. Homology regions of 105 bp located directly upstream of the stop codon, and 50 bp located 286 bp downstream of the stop codon within the 3’ UTR. The PAM was mutated such that glycine at position 452 was replaced with alanine. The same guide RNA and downstream homology was utilised in the knockout strategy. New upstream homology was selected, consisting of 50 bp located 271 bp upstream of the *COWP8* gene start codon.

The same gRNA and downstream homology for *Cp*COWP8 C-terminal tagging was used to generate *Δcowp8*. New upstream homology was selected, consisting of 50 bp located 271 bp upstream of the *CpCOWP8* start codon.

To generate COWP9-mScarlet-I, *CpCOWP9* (cgd6_210) was targeted with a guide RNA located 31 bp upstream of the stop codon. Upstream homology was already cloned into the tagging construct. Downstream homology of 50 bp located 266 bp after the stop codon.

### Excystation

Wild type *C. parvum* oocysts were incubated in 4% bleach on ice for 5 minutes, and then washed three times with PBS. Oocysts were incubated for 1 hour at 37°C in either 0.2 mM sodium deoxytaurocholate in PBS [31], or 0.5% (w/v) taurodeoxycholic acid in 0.25% trypsin in PBS [32] to induce excystation.

### Animal Ethics

Mice randomly assigned to cages at the time of weaning. Both sexes were used for experiments described here. All animal studies were ethically reviewed and carried out in accordance with Animals (Scientific Procedures) Act 1986 and the Dundee University Policy on the Care, Welfare, and Treatment of Animals. Regulated procedures on living animals were approved by the University’s Ethical Review Committee and carried out under the authority of project and personal licenses issued by the Home Office under the Animals (Scientific Procedures) Act 1986, as amended in 2012 (and in compliance with EU Directive EU/2010/63). The ERC has a general remit to develop and oversee policy on all aspects of the use of animals on University premises and is a subcommittee of the University Court, its highest governing body.

### Propagation of Transgenic Strains in Immunocompromised Mice

IFN-gamma KO mice (Jackson Laboratory #B6.129S7-IfngtmlTS/J, JAX 002287 bred at the University of Dundee; at least 8 weeks old) were given antibiotics in the drinking water (1 mg/ml ampicillin, 1 mg/ml streptomycin, 0.5 mg/ml vancomycin final concentrations) for 1 week prior to infection. Excysted sporozoites were transfected using a 4D Amaxa Nucleofector as previously described.[31] Mice were gavaged with saturated sodium bicarbonate 5 min prior to infection by gavage with transfected sporozoites. Both sexes used to generate new strains of transgenic *Cryptosporidium* (2 mice). Paromomycin (16 g/L) was added to the drinking water for the duration of the infection, or as indicated.

To passage transgenic strains, mice were gavaged with faecal slurry or purified oocysts. Mice were treated with selection agents as indicated. Faecal samples were collected post infection as indicated and were pooled from the cage of mice.

### Faecal Sample Analysis

NanoLuciferase assay was performed on 20 mg of homogenised faecal sample (in 1mL of lysis buffer). 25 µL of NanoLuciferase substrate-lysis buffer mixture was added to 100 µL of faecal lysate (GloMax plate reader, Promega). Positive threshold for NanoLuciferase signal is RLU >1000. Oocysts were purified from faecal samples using sucrose and cesium chloride flotations as previously described.

DNA was extracted from either ∼100 mg faecal material or purified oocysts. Samples were subjected to minimum of five freeze thaw cycles and then DNA was extracted using either a Zymo Quick-DNA Faecal/Soil Microbe DNA Miniprep Kit (faecal sample) or a Qiagen DNeasy Blood & Tissue Kit (oocysts). DNA was amplified by PCR using primers specific to the 5’ and 3’ regions of the repair cassette integration site. For knockout mutants, a third set of primers designed to amplify within the targeted genes ORF were also used. *α-tubulin* served as the positive control.

### Whole genome sequencing and analysis

In the case of *Δcowp8*, DNA was extracted from 10 million purified oocysts and sequenced and analysed by Glasgow Polyomics. Briefly, library was prepared using NEBNext UItra II FS DNA kit, which uses enzymatic fragmentation. Library was then sequenced 2*100bp to an average of >5 million reads on an Illumina NextSeq2000. The FASTQ files forward and reverse paired-end reads of COWP8 KO sample were aligned to the reference genome v68 of *C. parvum* clone IOWAII downloaded from CriptoDB[33] using Bowtie2 version 2.3.5[34], with the ‘very-sensitive-local’ pre-set alignment option. The alignments were converted to BAM format, reference sorted and indexed with SAMtools version 1.9.[35] The genome coverage of the aligned reads was extracted from the BAM files using bedtools version v2.29.0[36] with the -pc option for pair end data and -bg option to output a bedGraph file.

The GFF annotation file for *C. parvum* clone IOWAII was downloaded from CriptDB and converted to GTF with gffread (https://github.com/gpertea/gffread). The bedGraph file and GTF file were visualized with the svist4get python package version 1.2.24.[37]

### Fluorescence Microscopy

#### Live oocysts

Live oocysts were bleached, washed, and resuspended in a 1:1 mix of matrigel (Corning):PBS. Oocysts were loaded onto a µ-Slide Angiogenesis slide (Ibidi, catalog #81501) and allowed to settle overnight at 4°C. Oocyst:matrigel mixture was incubated at 37°C for 15 minutes immediately before imaging to solidify matrigel and immobilize oocysts.

#### Excysted oocysts

One million oocysts were excysted as described above, and then washed once in PBS, and fixed for 10 min in 4 % paraformaldehyde in PBS. After a PBS wash, samples were permeabilised with 0.25% Triton X-100 in PBS for 10 min. After a PBS wash, samples were blocked using 3% bovine serum albumin (BSA) (w/v) in PBS for at least 30 minutes. Oocysts were labelled with primary antibody (antibodies prepared in 3% BSA in PBS) for 1 hour. Samples were washed with PBS and stained with secondary antibody and Hoechst 33342 (Thermo Scientific) for 1 hour. After a final wash, oocysts were resuspended in 20 μL of either ProLong Diamond Antifade Mountant or ProLong Glass Antifade Mountant (Invitrogen). Resuspended oocysts were pipetted between a glass slide and coverslip (ThorLabs, #1.5H thickness, 22 × 22 mm).

#### Intracellular Stages

HCT-8 cells (ATCC catalogue #CCL-244; RRID: CVCL_2478) were cultured on sterile 12mm glass coverslips in a 24-well plate. Once cells reached a confluency between 40-60%, bleached and washed oocysts were added. Cultures were fixed at the indicated time points with 3 % paraformaldehyde, quenched with 125 mM glycine and permeabilized with 0.05 % saponin in PBS (10 minutes). Samples were labelled as described for excysted oocysts.

Intestinal tissue was harvested at peak infection. The distal ileum was flushed with PBS and fixed overnight, gently rocking in 4% (w/v) paraformaldehyde at 4°C. Tissue was dehydrated in 30% (w/v) sucrose in PBS overnight at 4°C and embedded in OCT compound (Agar Scientific) and cryosectioned (10 µm thick, Leica CM1850 Cryostat). Sections were dried overnight onto Thermo Scientific SuperFrost Plus Adhesion slides. Samples were stained with Hoechst 33342 (Thermo Scientific) and Phalloidin-647 (Abcam, catalog #ab176759) and mounted using ProLong Glass Antifade (Invitrogen).

#### Fluorescence Imaging

All antibodies and dyes used are described in **Supplemental Table 3**. All images collected at the Dundee Imaging facility; the Open Microscope Environment (OMERO) was utilised for image management. Z-stacked images of tissue sections were acquired using a DeltaVision Widefield Deconvolution microscope or a Zeiss LSM 880 (optionally with Airyscan detector) as indicated.

### SEM and TEM microscopy

For routine transmission electron microscopy and immunoEM, four million oocysts (wild type or COWP8-mNeon) were bleached and washed three times in PBS (excluding *Δcowp8* oocysts). All oocyst samples were fixed overnight at 4 °C in EM-grade 4% paraformaldehyde and either 2.5% (for standard TEM) or 0.2% (for immunogold TEM) glutaraldehyde in PBS. Fixed parasites were sent for processing and imaging at the Cellular Analysis Facility, University of Glasgow.

For routine TEM, oocysts were washed in 0.1 M phosphate buffer, before being post-fixed in 1% OsO_4_ and 1.25% potassium ferrocyanide (vol:vol) for 1 hour on ice in darkness. Oocysts were then washed in distilled water following contrast en bloc with 0.5 % aqueous uranyl acetate for 1 hour at room temperature in darkness. Samples were then dehydrated in an ascending series of acetone and embedded in epoxy resin. Ultra-thin sections (60 nm) were cut and collected onto formvar-coated copper grids and contrasted with 4% uranyl acetate prior imaging using a JEOL 1200 EX transmission electron microscope, operating at 80kV.

For immunoEM, after several washes in the cacodylate 0.1M buffer, the samples were dehydrated in ascending ethanol series and embedded in LR White resin (Agar Scientific). Ultrathin sections (60 nm thick) were obtained using an ultramicrotome (Leica Microsystems). The sections were collected on formvar-coated nickel grids and then blocked in PBS containing 3% bovine serum albumin for 1 hour. After this time, they were incubated in the presence of primary antibody (either a monoclonal mouse anti-c-myc (ProteinTech) or a monoclonal rat anti-HA (SigmaAldrich)). Then they were washed several times in blocking buffer and incubated with 15 nm gold-conjugated Protein A (Aurion). The grids were washed several times in the blocking buffer, dried and contrasted with 4% uranyl acetate, and observed using a JEOL 1200 EX transmission electron microscope operating at 80kV.

For scanning electron microscopy two million oocysts were bleached and washed three times in PBS. Oocysts were fixed at 4 °C overnight in EM-grade 4% paraformaldehyde and 2.5% glutaraldehyde in PBS. Fixed oocysts were washed in 0.1 M phosphate buffer, then left on top of poly-l-lysine coated glass slides for 20 min to allow oocysts to adhere. Adhered oocysts were washed in 0.1 M phosphate buffer, before being dehydrated in an ascending series of ethanol concentrations, and finally reaching critical point dehydration with a Autosamdri-815 machine (Tousimis). Dried coverslips were coated with gold/palladium (10 nm thick layer) and imaged in a JEOL IT-100 scanning electron microscope.

### *In vivo* phenotyping experiments

25,000 oocysts were resuspended in PBS and exposed to 60°C in an Eppendorf ThermoMixer. A temperature of 60 °C was chosen, as this temperature is at the lower end of reported temperatures known to heat inactivate *Cryptosporidium*.[38–40] Samples were then stored at 4°C overnight.

50,000 oocysts were resuspended in either 0% or 8% undiluted bleach solution and incubated on ice for 5 minutes. Oocysts were washed in PBS three times and then stored at 4°C overnight.

Age and sex matched interferon-gamma knockout mice (4 mice were cage) were pre-treated prior to infection with 5,000 oocysts per strain by oral gavage.

### Nanoindentation

Petri dishes (35 mm × 10 mm, Corning cat no. 430588) were treated with Corning Cell-Tak cell and tissue adhesive (cat no. CLS354240) the day before oocyst attachment. Cell-Tak was added to 0.1 M sodium bicarbonate solution, applied to 35 mm dish, and incubated for a minimum of 20 min at room temperature. Cell-Tak solution was aspirated and dish washed twice with sterile water. Dishes were left to air dry and stored in at 4°C.

One million *Δcowp8* and WT oocysts in PBS were applied to a dish and incubated overnight at 4 °C. Dishes were gently washed once with sterile PBS and stored in sterile PBS to prevent oocysts desiccation. Nanoindentation measurements and analysis were conducted at the Advanced Research Centre (ARC), University of Glasgow. Nanoindentation measurements were performed using a Chiaro Nanoindenter (Optics 11) mounted on top of an inverted phase contrast microscope (Evos XL Core, Thermofisher), following a previously described approach.[41] Measurements were performed at room temperature in PBS. Each sample was indented once with a spherical tip of 22 μm radius. Measurements were performed on oocysts adhered to three separates petri dishes, sampling a minimum of ten oocysts per plate (n=60-67) and the experiment was repeated twice (n=2).

The selected cantilever had a stiffness of 4.18 Newton metres (Nm–1). The collected curves were analysed using a custom Python code.[42] Curves were first aligned using a baseline detection method based on the histogram of the force signal[43] and the corresponding indentation was calculated for each curve. The analysis was performed using the Hertz model for a spherical indenter to fit the curves obtained.

## Proteomics

### Sample preparation and protein extraction

Samples of wild type oocysts and *Δcowp8* oocysts were prepared as two biological repeats (with two technical repeats each) and analyzed using quantitative proteomics. 40 million oocysts per technical repeat were bleached, washed three times in PBS, and excysted at 37 °C in 0.2 mM sodium taurochlorate for 2 hours. Excysted material was resuspended in 25 µL lysis buffer (4% SDS, 10 mM DTT, 1x protease inhibitor cocktail, all prepared in water). Sample was subjected to five cycles of freeze thaw to excyst remaining oocysts. Samples were incubated at 56 °C for 1 hour while shaking (800 rpm) to solubilize proteins. Samples were centrifuged at 20,000xg for 40 minutes at 4 °C. Supernantant was removed and TCEP (*tris(2-carboxyethyl)phosphine*) added to a final concentration of 25 mM and incubated at 37 °C for 10 minutes. Added IAA (iodoacetamide) to a final concentration of 25 mM and incubated at room temperature in the dark for 1 hour. Added 13% volume ice-cold TCA and store at -20 C. Sample was centrifuged at 16,000 xg for 4 °C for 5 minutes. Supernatant was removed and pellet was washed with 0.5 ml cold acetone. Acetone wash repeated a total of five times and then the sample was air dried. Protein extract was resuspended in 400 µl TEAB, sonicated to resuspend the material. Sample was digested with Trypsin/LysC (Promega) overnight at 37 °C with shaking, and then dried.

### TMT labelling and high pH reverse phase fractionation

Tryptic peptides (11.6µg, from each sample) were dissolved in 100 µl of 150 mM TEAB. TMT labelling was performed according to the manufacturer’s instructions (Thermo-Fisher Scientific). The different TMT-10 plex labels (0.8mg) (Thermo Fisher Scientific) were dissolved in 41µL of anhydrous acetonitrile, and each label is added to a different sample. The mixture was incubated for 1 hour at room temperature, an equivalent of 0.75 µg of peptides from each sample were mixed with 20 µl 1% formic acid and used to check labelling efficiency. The remaining samples were kept at -80°c, until the labelling efficiency was checked. Samples were pooled, desalted, and dried in a speed-vac at 30°C. Samples were re-dissolved in 200 µl ammonium formate (10mM, pH 9.5) and peptides were fractionated using High pH RP Chromatography. A C18 Column from Waters (XBridge peptide BEH, 130Å, 3.5 µm 2.1 X 150 mm, Waters, Ireland) with a guard column (XBridge, C18, 3.5 µm, 2.1X10mm, Waters) were used on an Ultimate 3000 HPLC (Thermo-Scientific). Buffers A and B used for fractionation consist, respectively, of (A) 10 mM ammonium formate in milliQ water pH 9.5 and (B) 10 mM ammonium formate, pH 9.5 in 90% acetonitrile. Fractions were collected using a WPS-3000FC auto-sampler (Thermo-Scientific) at 1minute intervals. Column and guard column were equilibrated with 2% Buffer B for twenty minutes at a constant flow rate of 0.2ml/min. Fractionation of TMT labelled peptides was performed as fellows; 190 µl aliquot were injected onto the column, and the separation gradient was started 1 minute after the sample was loaded onto the column. Peptides were eluted from the column with a gradient of 2% Buffer B to 20% Buffer B in 6 minutes, then from 20% Buffer B to 45% Buffer B in 51 minutes and finally from 45% buffer B to 100% Buffer B within 1 min. The Column was washed for 15 minutes in 100% Buffer B. The fraction collection started 1 minute after injection and stopped after 80 minutes (total 80 fractions, 200µl each). Formic acid (30 µl of 10% stock) was added to each fraction and concatenated in groups of 20 fractions.

### LC-MS analysis

Analysis of peptides was performed on a Q-exactive-HF (Thermo Scientific) mass spectrometer coupled with a Dionex Ultimate 3000 RS (Thermo Scientific). LC buffers were the following: buffer A (0.1% formic acid in Milli-Q water (v/v), buffer B (80% acetonitrile and 0.1% formic acid in Milli-Q water (v/v) and loading buffer (0.1% TFA).

Peptides from each fraction were resuspended in 50 µl 1% formic acid and aliquots of 5 μL were loaded at 10 μL/min onto a trap column (100 μm × 2 cm, PepMap nanoViper C18 column, 5 μm, 100 Å, Thermo Scientific) equilibrated in 0.1% TFA. The trap column was washed for 5 min at the same flow rate with 0.1% TFA and then switched in-line with a Thermo Scientific, resolving C18 column (75 μm × 50 cm, PepMap RSLC C18 column, 2 μm, 100 Å) equilibrated in 5% buffer B for 17 min. The peptides were eluted from the column at a constant flow rate of 300 nl/min with a linear gradient from 5% buffer B (for Fractions 1-10, 7% for Fractions 11-20) to 35% buffer B in 125 min, and then from 35% buffer B to 98% buffer B in 2 min. The column was then washed with 98% buffer B for 20 min and re-equilibrated in 5% or 7% buffer B for 17 min. The column was maintained at a constant temperature of 50°C.

Q-exactive HF was operated in data dependent positive ionisation mode. The source voltage was set to 2.85 Kv and the capillary temperature was 250°C. A scan cycle comprised MS1 scan (m/z range from 335-1600, with a maximum ion injection time of 50 ms, a resolution of 120 000 and automatic gain control (AGC) value of 3x10^6^) followed by 15 sequential dependant MS2 scans (resolution 60000) of the most intense ions fulfilling predefined selection criteria (AGC 1x10^5^, maximum ion injection time 200 ms, isolation window of 0.7 m/z, fixed first mass of 100 m/z, spectrum data type: centroid, intensity threshold 5 x 10^4^, exclusion of unassigned, singly and >6 charged precursors, peptide match preferred, exclude isotopes on, dynamic exclusion time of 45 s). The HCD collision energy was set to 32% of the normalized collision energy. Mass accuracy is checked before the start of samples analysis.

### Quantitative proteomics analysis of transgenic *Cryptosporidium*

Proteomic data were processed through MaxQuant software (version 2.4.10.0), leveraging its integrated Andromeda search engine.[44] The search database was specifically constructed for *Cryptosporidium parvum* Iowa II, with annotated protein sequences obtained from CryptoDB,[33] release 61. This was augmented with sequences for the reporter proteins mScarlet-I and NLuc-Neo. Additionally, a murine protein database, retrieved from Uniprot[45] on January 20, 2023, was concatenated with the *C. parvum* database to account for potential host protein contamination. The analysis encompassed both TMT 6-plex and TMT 10-plex labeling, processed in parallel within a single MaxQuant instance. Each TMT experiment was treated as a distinct analytical batch, with no normalization applied between them. Trypsin was designated as the proteolytic enzyme, with a specification for up to two missed cleavages per peptide allowed. Carbamidomethylation of cysteine residues was set as a constant post-translational modification (PTM), whereas N-terminal acetylation of proteins and methionine oxidation were configured as variable PTMs. Default settings were retained for all other parameters within MaxQuant, except for TMT label correction factors. These were adjusted according to the manufacturer’s instructions (Thermo Fisher Scientific) and are detailed within the MaxQuant parameter files. These files, alongside the raw data, have been submitted to the PRIDE database (PRIDE submission in progress). Data is available under Project accession: PXD050089.

Pre-normalization, we filtered the MaxQuant output to exclude potential confounders. Data were refined by removing entries solely identified by peptide modification sites (Only identified by site), entries marked as reverse database matches (Reverse), and proteins classified as potential contaminants (Potential contaminant). For our analysis of *Cryptosporidium parvum* COWP8 across wild-type (WT) and knockout (KO) specimens, we used a robust normalization technique tailored specifically for TMT proteomic data. Our experimental set consisted of one biological replicate with two technical replicates integrated within a TMT 6-plex configuration, along with an additional biological replicate paired with two technical replicates in a TMT 10-plex setup. To normalize the signal of the two TMT batches, we adopted a modified version of the Internal Reference Standard (IRS) normalization method, as initially delineated by Plubell et al.[46] The IRS normalization was applied by taking the raw mean intensity of each plex to serve as the reference channel. We computed the sum of the intensities for each TMT channel row-wise and then calculated the geometric mean of these sums to establish a stable reference point. This method was chosen to replace the reference channel utilized in,[46] which was not generated from our datasets. The scaling factors for normalization were ascertained by dividing this geometric mean by the sum intensities of each individual experiment. Subsequently, each plex underwent individual scaling to ensure uniformity in signal intensities. Upon normalizing the data, we employed Principal Component Analysis (PCA) to verify the efficacy of the IRS process between the TMT 6-plex and 10-plex setups. The differential expression analysis was performed with the limma package[47] using the WT samples versus the KO samples, with log2 values. FDR values were computed with the toptable function in limma. The output table of the analysis is available as attached csv file (**Supplemental Table 4**).

### Statistics

Graphs prepared using GraphPad Prism version 10.2.1.

## Acknowledgements

Thanks to Beatrice L. Colon, Flora C Caldwell, Rachel Chalmers, Susan Wyllie, David Horn, Marcus Lee and the Lee lab for helpful discussion. We would like to acknowledge the University of Dundee School of Life Sciences Resource Unit, Dundee FingerPrints Proteomics Facility, Dundee Imaging Facility, and Dundee Flow Cytometry and Cell Sorting Facility (supported by the Wellcome Trust, 097418/Z/11/Z) and University of Glasgow Polyomics.

