## Supplemental Figures and Tables for "*Cryptosporidium* Oocyst Wall Proteins are true oocyst wall proteins, with COWP8 functioning to hold the inner and outer layers of the oocyst wall together"

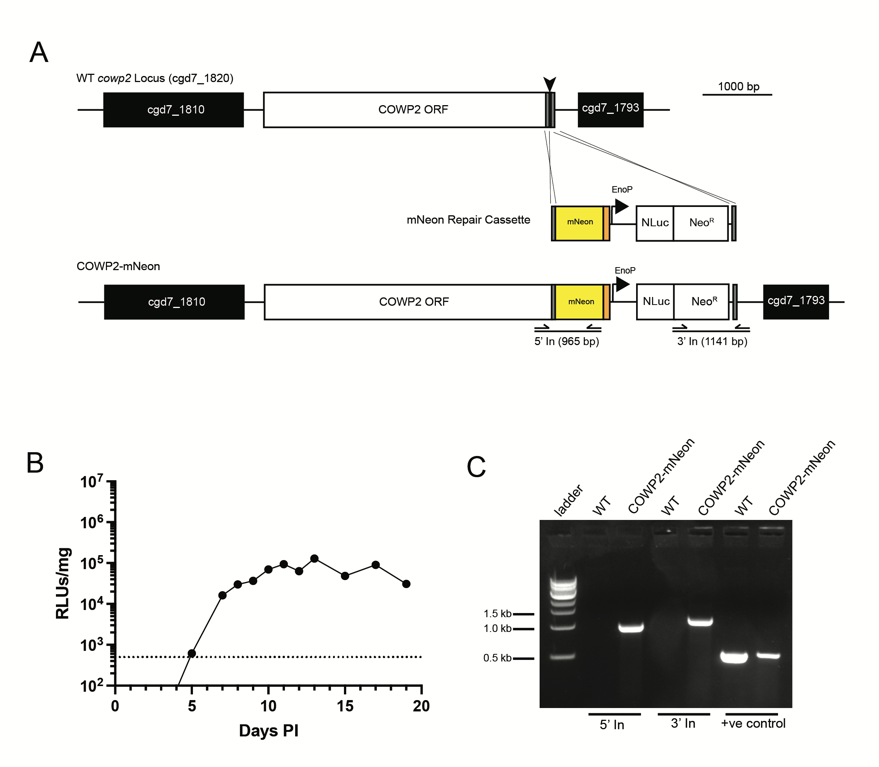

**Supplemental Figure 1. Generation of COWP2 tagged strain.**

**A**) gRNA (black arrow) and regions of 50 bp of homology (gray) were used to target the C-terminus of COWP2 (cgd7_1820) for fusion with mNeon (mNeon in yellow and 3xHA in orange). Strain includes NanoLuciferase-Neomycin resistance fusion protein (NLuc-Neo^R^) expressed by the constitutive *CpEnolase* promoter. **B**) Infection level of mice as measured by faecal NLuc, limit of detection at 500 RLU/mg, dotted line. Average and SD of three technical replicates of one biological replicate. The first passage of COWP2-mNeon was well above the limit of detection. **C**) PCR with primer pairs indicated in (**A**) was performed using genomic DNA extracted from wild type and transgenic strains. PCR products confirm correct integration of (**C**) mNeon at the C-terminus of COWP2.

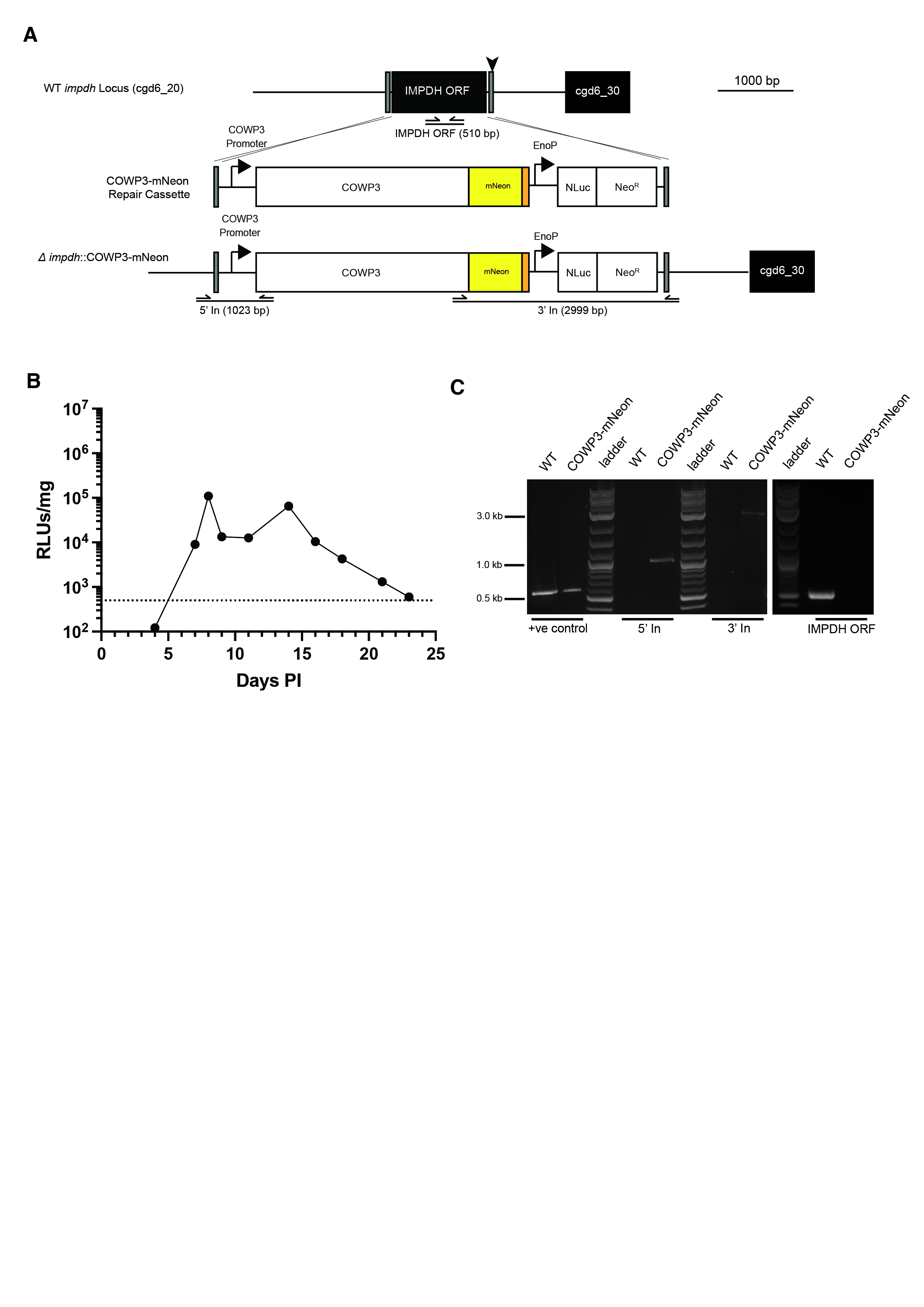

**Supplemental Figure 2. Expression of COWP3-mNeon tagged strain at *CpIMPDH* locus.**

**A**) Promoter and full open reading frame (ORF) of COWP3 cloned with C-terminal mNeon (same construct for endogenous C-terminal tagging as reported in **Supplemental Figure 1A**) to generate COWP3-mNeon repair cassette. This was targeted for integration at *CpIMPDH* locus (cgd6_20) using gRNA (black arrow) and regions of 50 bp of homology (gray). **B**) Infection level of mice as measured by faecal NLuc, limit of detection at 500 RLU/mg, dotted line. Average and SD of three technical replicates of one biological replicate. The first passage of *∆impdh*::COWP3-mNeon was well above the limit of detection. **C**) PCR with primer pairs indicated in (**A**) was performed using genomic DNA extracted from wild type and *∆impdh*::COWP3-mNeon.

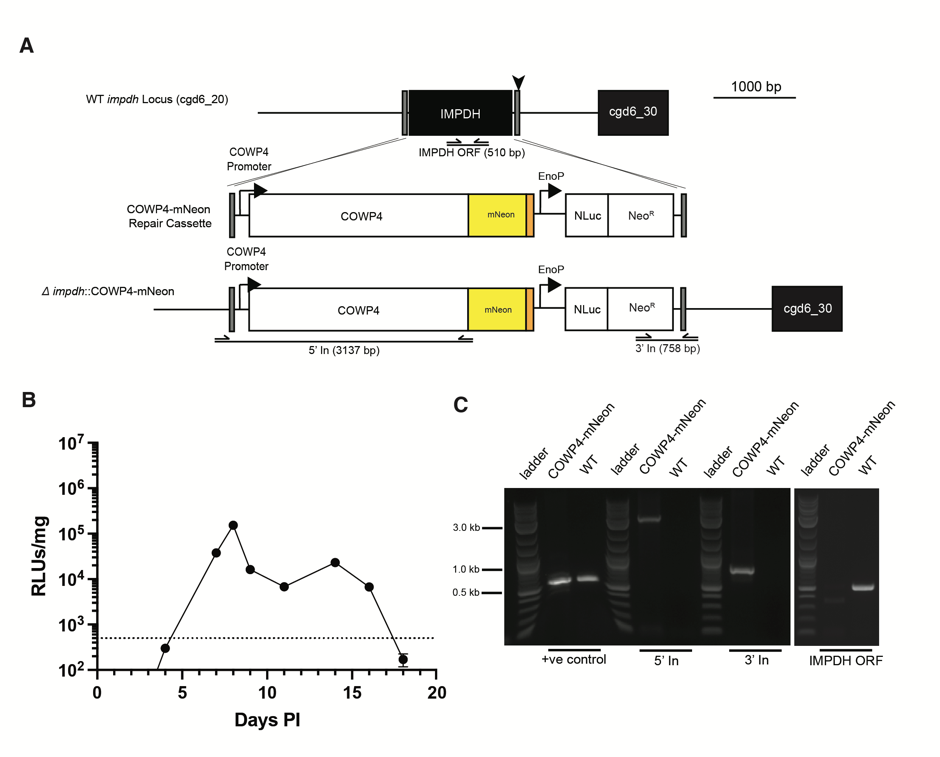

**Supplemental Figure 3. Expression of COWP4-mNeon tagged strain at *CpIMPDH* locus.**

**A**) Promoter and ORF of *COWP4* cloned with C-terminal mNeon to generate COWP4-mNeon repair cassette. This was targeted for integration at *CpIMPDH* locus (cgd6_20) using gRNA (black arrow) and regions of 50 bp of homology (gray). **B**) Infection level of mice as measured by faecal NLuc, limit of detection at 500 RLU/mg, dotted line. Average and SD of three technical replicates of one biological replicate. The first passage of *∆impdh*::COWP4-mNeon was well above the limit of detection. **C**) PCR with primer pairs indicated in (**A**) was performed using genomic DNA extracted from wild type and *∆impdh*::COWP4-mNeon.

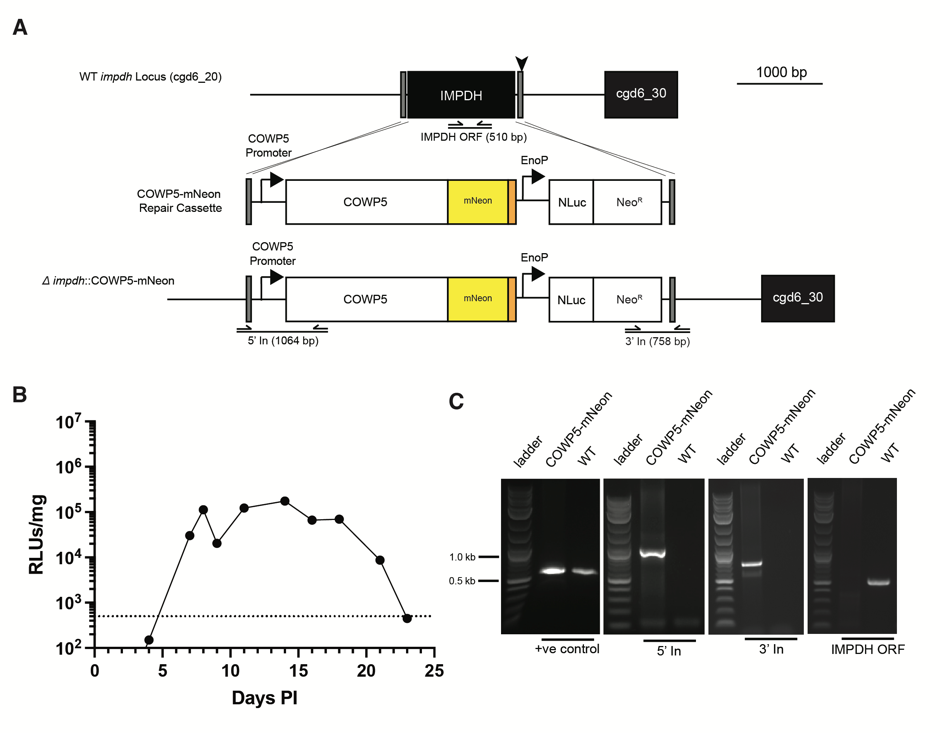

**Supplemental Figure 4. Expression of COWP5-mNeon tagged strain at *CpIMPDH* locus.**

**A**) Promoter and ORF of *COWP5* cloned with C-terminal mNeon to generate COWP5-mNeon repair cassette. This was targeted for integration at *CpIMPDH* locus (cgd6_20) using gRNA (black arrow) and regions of 50 bp of homology (gray). **B**) Infection level of mice as measured by faecal NLuc, limit of detection at 500 RLU/mg, dotted line. Average and SD of three technical replicates of one biological replicate. The first passage of *∆impdh*::COWP5-mNeon was well above the limit of detection. **C**) PCR with primer pairs indicated in (**A**) was performed using genomic DNA extracted from wild type and *∆impdh*::COWP5-mNeon.

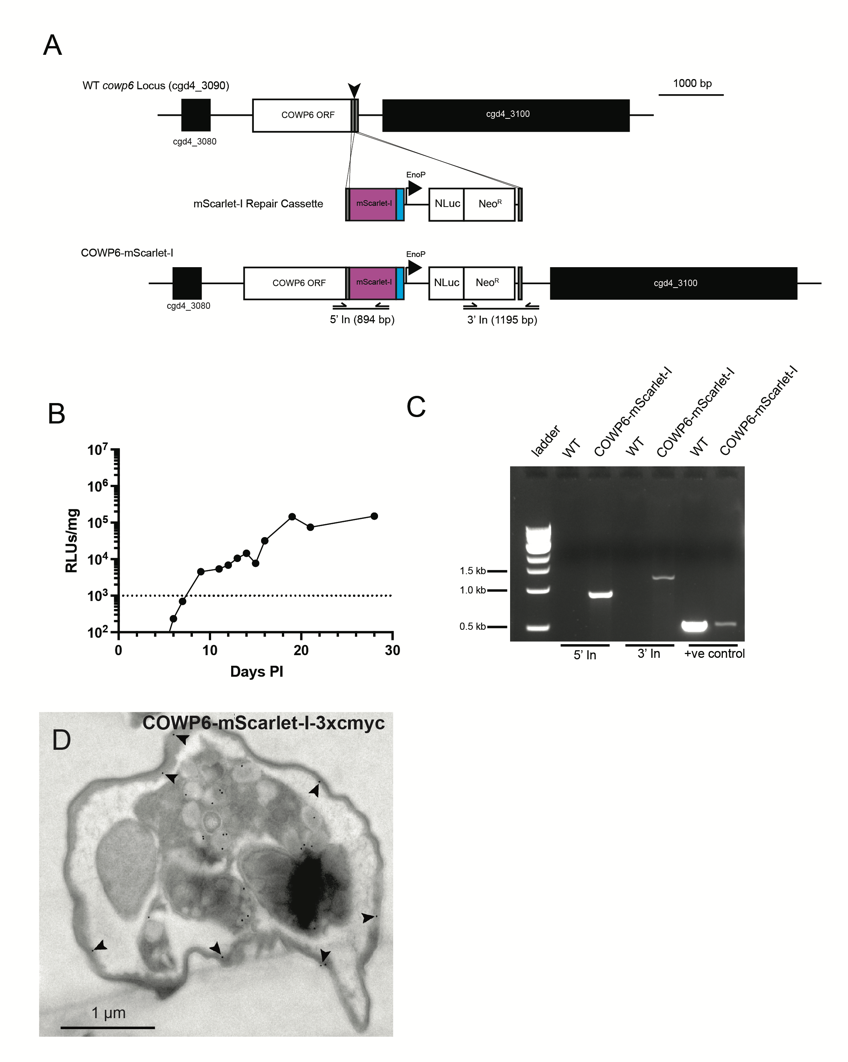

**Supplemental Figure 5. Generation of COWP6-mScarlet-I strain.**

**A**) Strategy to target the C-terminus of COWP6 (cgd4_3090) for fusion with the mScarlet-I-3xmyc Repair Cassette (mScarlet in magenta and 3xmyc in blue). NanoLuciferase-Neomycin resistance fusion protein (NLuc-Neo^R^) expressed by the constitutive Cp*Enolase* promoter. gRNA (black arrow) and regions of 50 bp of homology (gray). **B**) Infection level of mice as measured by faecal NLuc, limit of detection at 500 RLU/mg, dotted line. Average and SD of three technical replicates of one biological replicate. The first passage of COWP6-mScarlet-I (black circles) was well above the limit of detection. **C**) PCR with primer pairs indicated in (**A**) was performed using genomic DNA extracted from wild type and COWP6-mScarlet-I. **D**) Immunoelectron microscopy of COWP6-mScarlet-I confirm localization to the inner layer of the oocyst wall. Representative image shown.

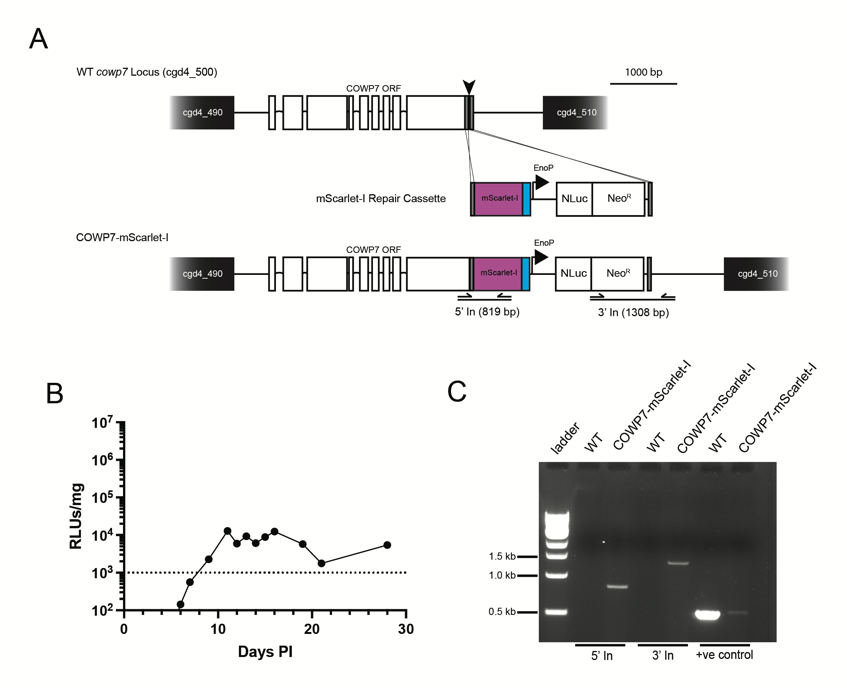

**Supplemental Figure 6. Generation of COWP7-mScarlet-I strain.**

**A**) Strategy to target the C-terminus of COWP7 (cgd4_500) for fusion with the mScarlet-I-3xmyc Repair Cassette (same construct for endogenous C-terminal tagging as reported in **Supplemental Figure 5A**). COWP7 is predicted to contain 8 introns; exons indicated by white boxes. NanoLuciferase-Neomycin resistance fusion protein (NLuc-Neo^R^) expressed by the constitutive Cp*Enolase* promoter. gRNA (black arrow) and regions of 50 bp of homology (gray). **B**) Infection level of mice as measured by faecal NLuc, limit of detection at 500 RLU/mg, dotted line. Average and SD of three technical replicates of one biological replicate. The first passage of COWP7-mScarlet-I (black circles) was well above the limit of detection. **C**) PCR with primer pairs indicated in (**A**) was performed using genomic DNA extracted from wild type and COWP7-mScarlet-I.

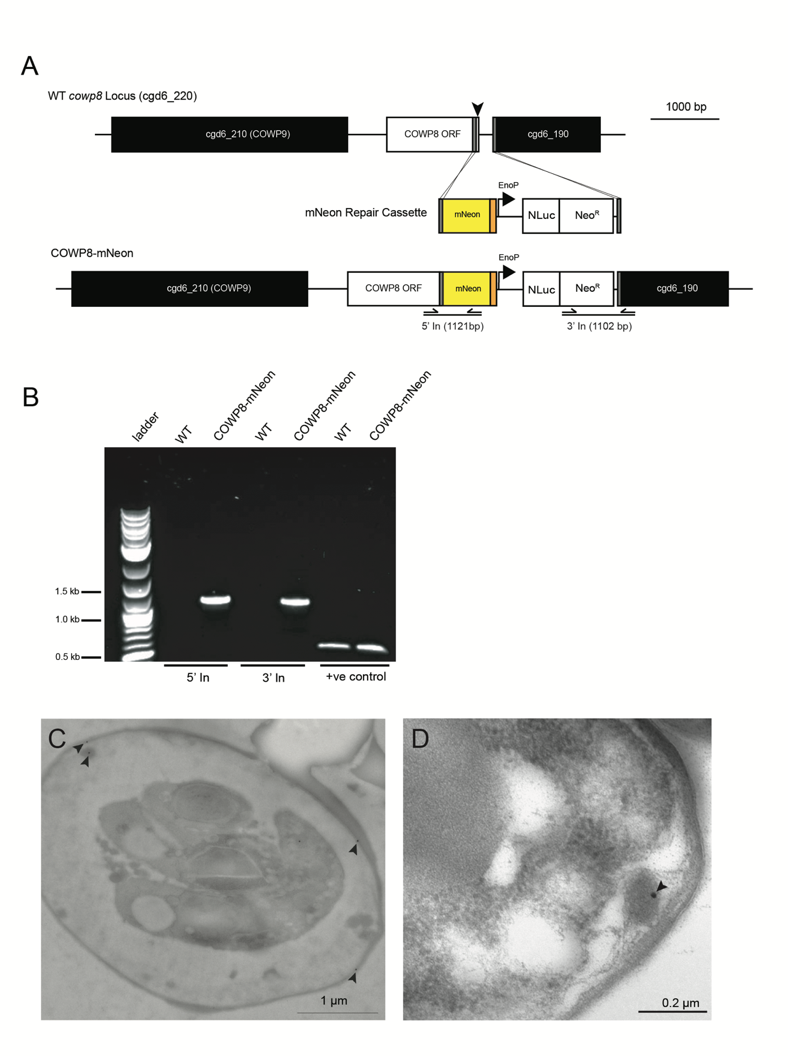

**Supplemental Figure 7. Generation of COWP8-mNeon strain.**

**A**) Strategy to target the C-terminus of COWP8 (cgd6_220) for fusion with the mNeon-3xHA Repair Cassette (mNeon in yellow and 3xHA in orange). NanoLuciferase-Neomycin resistance fusion protein (NLuc-Neo^R^) expressed by the constitutive Cp*Enolase* promoter. gRNA (black arrow) and regions of 50 bp of homology (gray). Note that neighbouring gene upstream of COWP8 is COWP9, illustrated in black. Mouse infections reported in **Figure 3A-F**. **B**) PCR with primer pairs indicated in (**A**) was performed using genomic DNA extracted from wild type and COWP8-mNeon. **C**) Immunoelectron microscopy of COWP8-mNeon confirm localization to the inner layer of the oocyst wall. Representative image shown. **D**) A second localisation in globular-type structures (globule does not appear to be membrane bound) in the space between the sporozoite membrane and inside of the oocyst wall was observed to be positive for COWP8 localisation by Immunoelectron microscopy. Representative image shown.

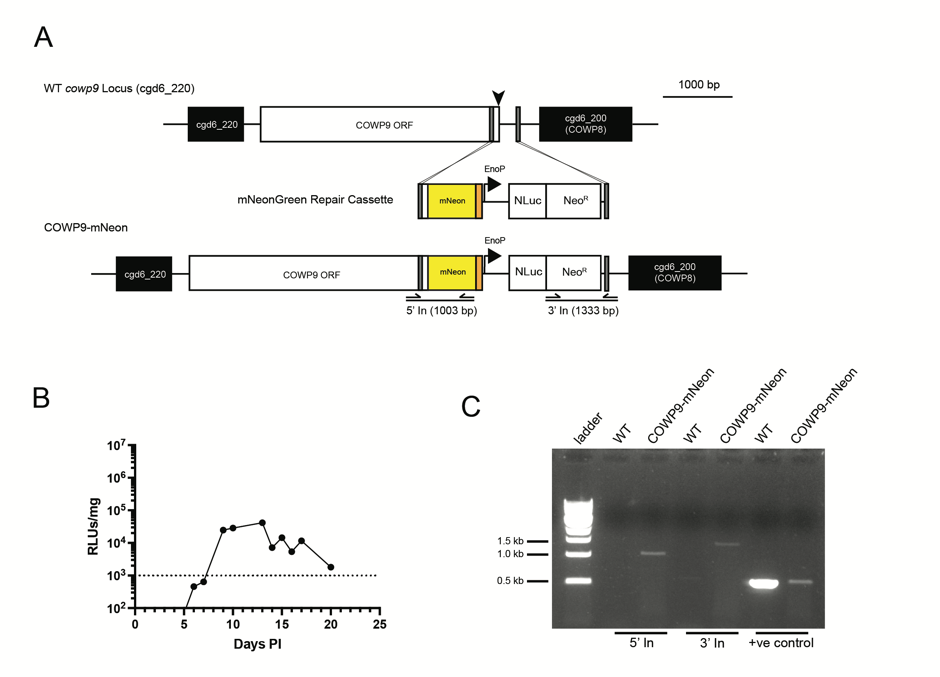

**Supplemental Figure 8. Generation of COWP9-mNeon strain.**

**A**) Strategy to target the C-terminus of COWP9 (cgd6_230) for fusion with the mNeon-3xHA Repair Cassette (mNeon in yellow and 3xHA in orange). NanoLuciferase-Neomycin resistance fusion protein (NLuc-Neo^R^) expressed by the constitutive Cp*Enolase* promoter. gRNA (black arrow) and regions of 50 bp of homology (gray). Note that neighboring gene downstream of COWP9 is CpCOWP8, illustrated in black. **B**) Infection level of mice as measured by fecal NLuc, limit of detection at 500 RLU/mg, dotted line. Average and SD of three technical replicates of one biological replicate. The first passage of COWP9-mScarlet-I (black circles) was well above the limit of detection. **C**) PCR with primer pairs indicated in (**A**) was performed using genomic DNA extracted from wild type and COWP9-mNeon.

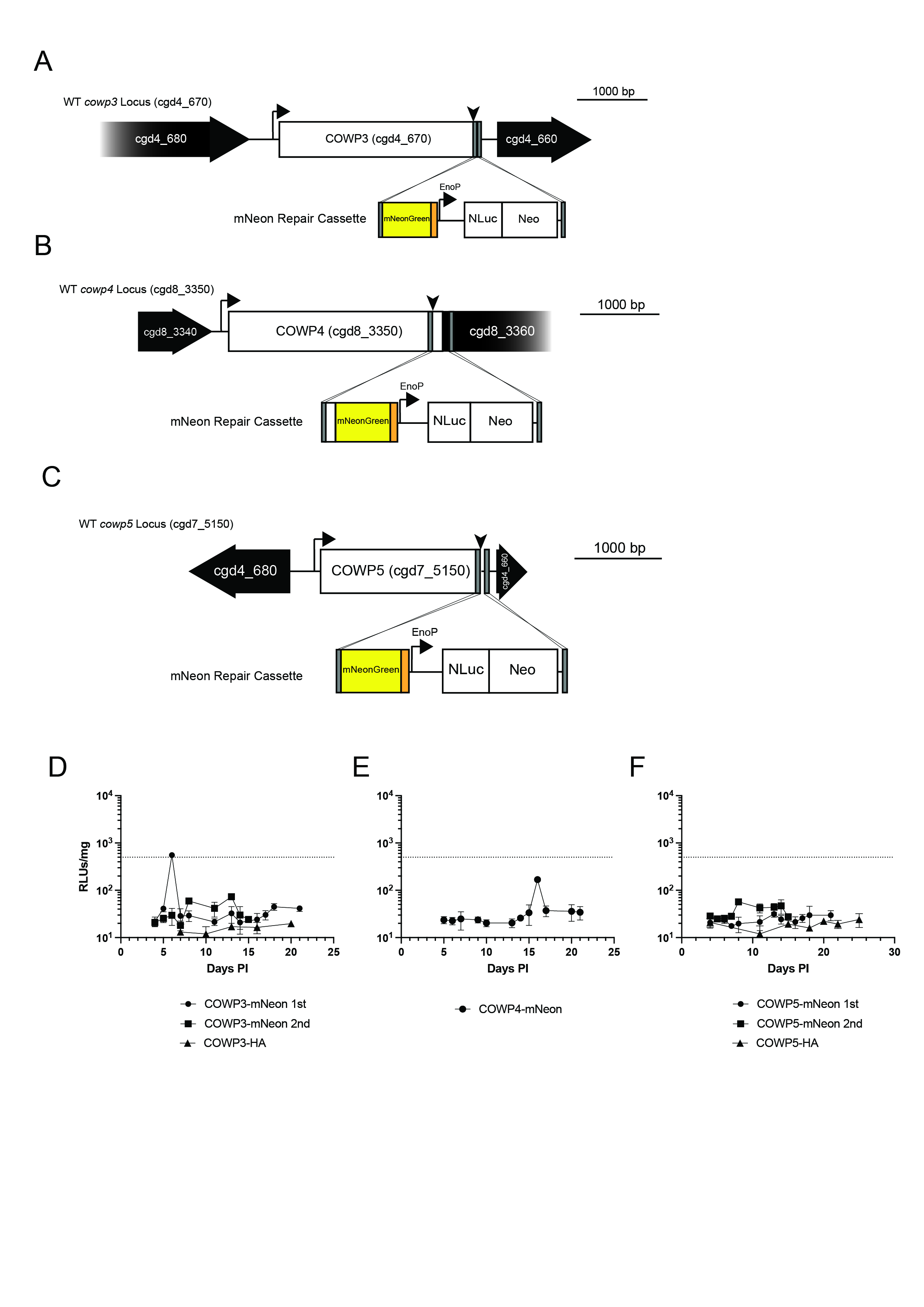

**Supplemental Figure 9. COWP3-5 cannot be endogenously C-terminally tagged.**

**A**) Strategy to target the C-terminus of COWP3 (cgd4_680) for fusion with the full mNeon-3xHA Repair Cassette, or the simplified 3xHA tag alone. Both strains include NanoLuciferase-Neomycin resistance fusion protein (NLuc-Neo^R^) expressed by the constitutive Cp*Enolase* promoter. gRNA (black arrow) and regions of 50 bp of homology (gray). Similar strategies designed for **B**) COWP4 (cgd8_3350) and **C**) COWP5 (cgd7_5150). Neighboring gene very near the C-terminus of each COWP is illustrated in black. **D-F**) Attempts to generate these mutants was unsuccessful as measured by fecal Nluc from infected mice; limit of detection at 500 RLU/mg, dotted line. Average and SD of three technical replicates of one biological replicate. **D**) COWP3 tagging was attempted twice with mNeon Repair cassette and once with simplified 3xHA tag. **E**) COWP4 tagging was attempted once with simplified 3xHA tag. **F**) COWP5 tagging was attempted twice with mNeon Repair cassette and once with simplified 3xHA tag.

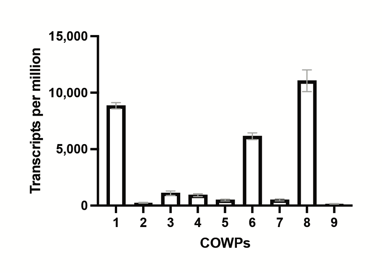

**Supplemental Figure 10. Expression of COWPs as determined from bulk RNASequencing of macrogamont life cycle stages.**

Transcript level of members of the COWP family from “female *in vivo*” sample from [22] as published on CryptoDB.org.[48]

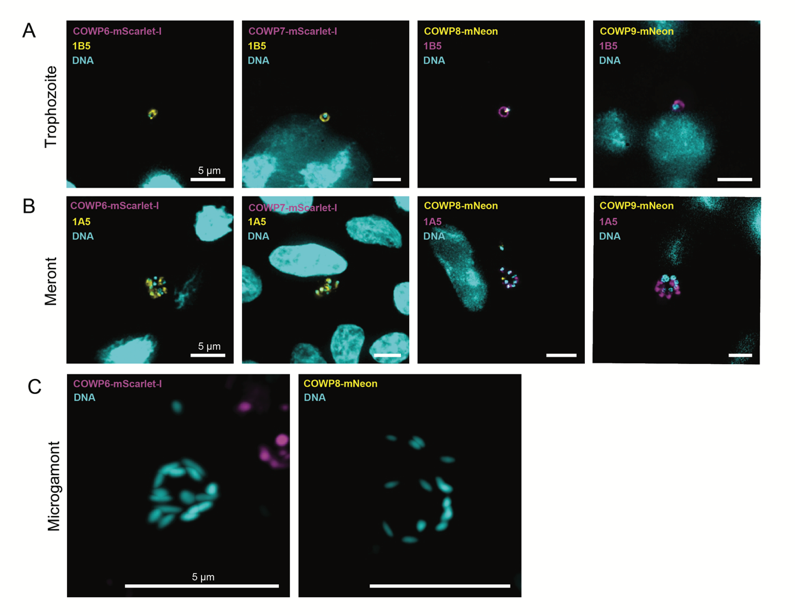

**Supplemental Figure 11. COWP6 and COWP8 are not expressed in asexual stages or microgamonts.**

Fluorescence microscopy of HCT-8 cells co-cultured individually with transgenic strains for (**A**) 12 hours or (**B**) 48 hours and fixed and processed for imaging. Single nuclei (DAPI, cyan) and staining with 1B5 (asexual marker, Sibley Lab Washington University, yellow or magenta as indicated) indicate trophozoite life cycle stage. Eight nuclei (DAPI, cyan) and staining with 1A5 (asexual marker, Sibley Lab Washington University, yellow or magenta as indicated) indicate meront life cycle stage. Images collected on a widefield epifluorescence microscope; representative images shown. **C**) Mice were culled at peak infection (faecal NLuc RLU/mg > 500,000) and processed for histology and immunofluorescence. Sixteen nuclei (DAPI, cyan), bullet shape and pattern as previously described used to categorize male parasites.[49] Super resolution images collected on a Zeiss LSM880 Airyscan microscope, airyscan mode.

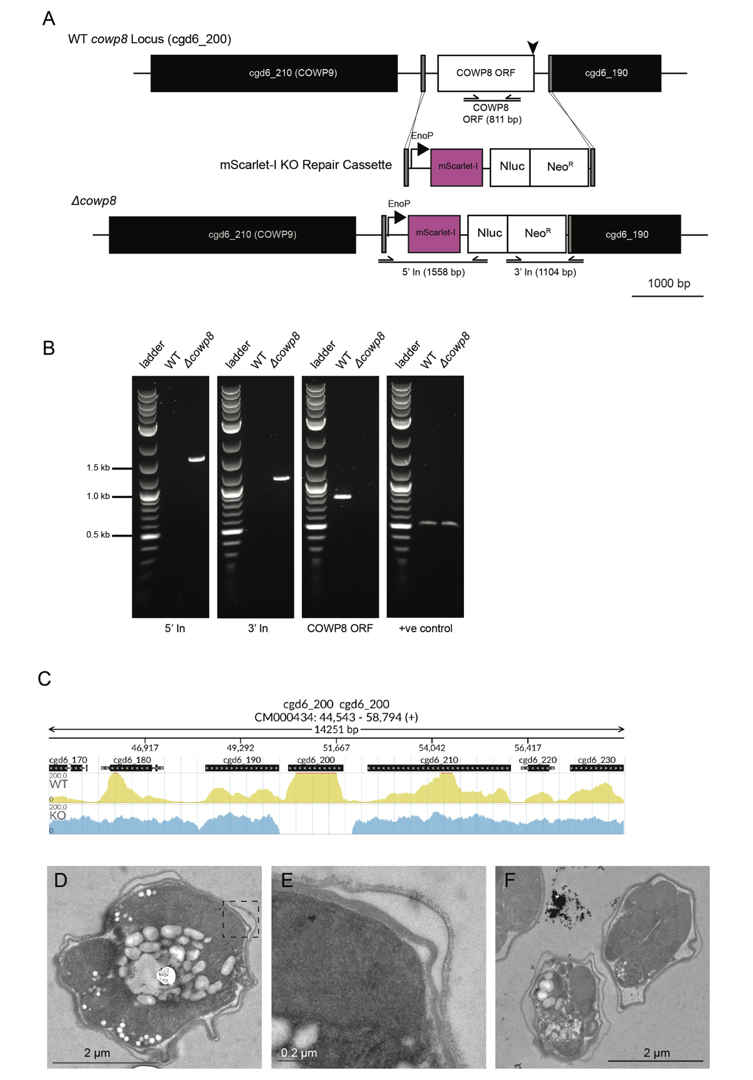

**Supplemental Figure 12. Generation of *∆cowp8*** **strain.**

**A**) Strategy to replace the full open reading frame of COWP8 (cgd6_220) with a repair cassette where mScarlet-I and NanoLuciferase-Neomycin resistance fusion protein (NLuc-Neo^R^) is expressed by the constitutive Cp*Enolase* promoter. The same gRNA (black arrow) that was used to target the C-terminus for florescent fusion and the same downstream region of 50 bp of homology (gray) was used to target the gene for deletion (**Supplemental Figure 7A**). Mouse infections reported in **Figure 3**. **B**) PCR with primer pairs indicated in (**A**) was performed using genomic DNA extracted from wild type and *∆cowp8.* **C**) Whole genome sequencing of *∆cowp8* confirms gene deletion. Genome coverage. For the COWP8 gene (cgd6_200) the figure shows the genome coverage for the KO (*∆cowp8*) sample. The top of the figure shows an ideogram of the gene structures. **D**) Additional representative images of *∆cowp8* oocyst, **E**) inset of D, and **F**) additional representative images of *∆cowp8* oocysts.

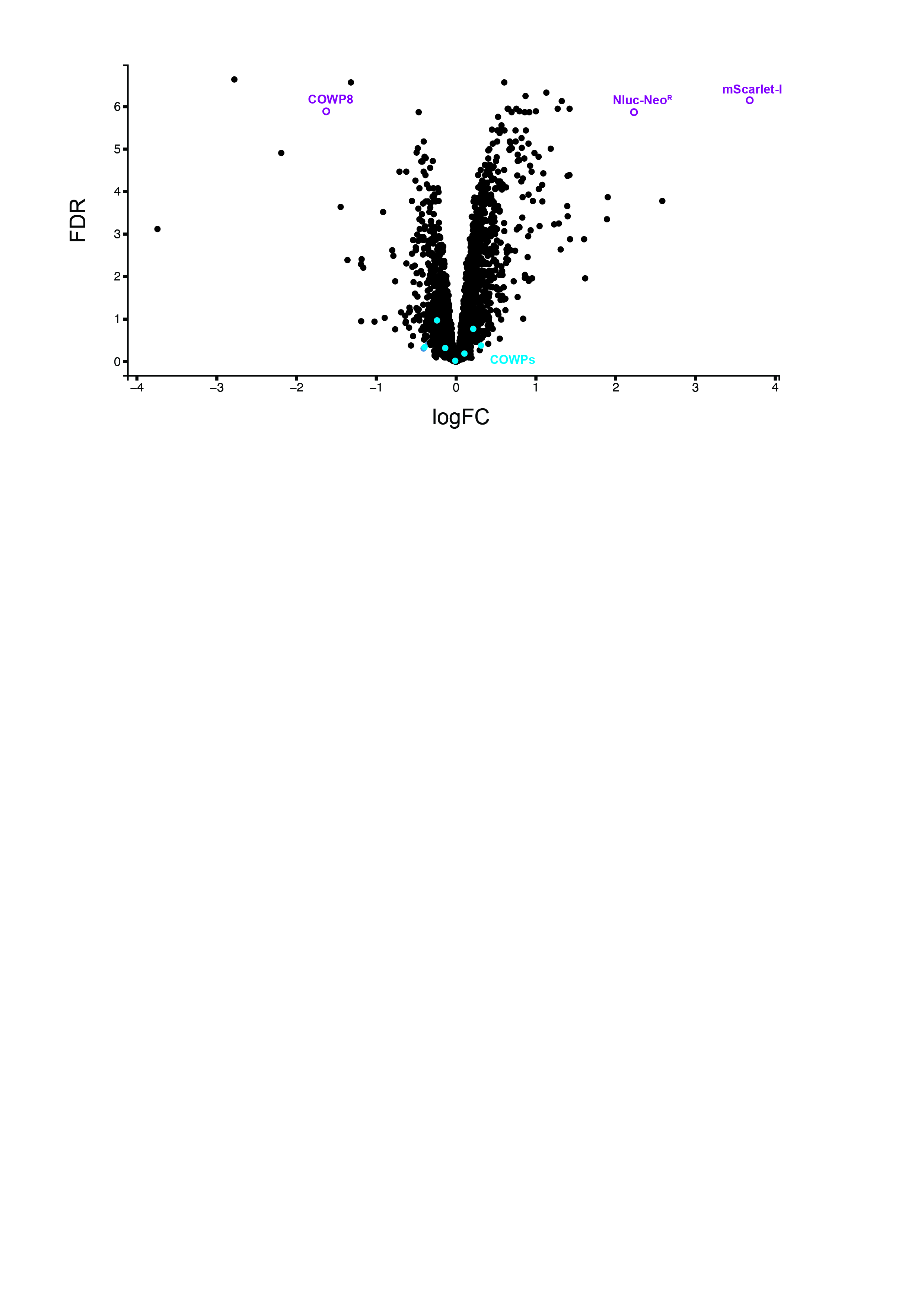

**Supplemental Figure 13. Loss of COWP8 does not change expression of other COWPs.**

Quantitative proteomics of wild type vs *∆cowp8* plotted as fold change (logFC) and false discovery rate (FDR). Protein extracted from excysted *Cryptosporidium* oocysts and samples labelled with tandem mass tags. COWPs indicated in blue, *∆cowp8* reporter proteins in purple. 2 biological replicates, each with 2 technical replicates. See **Supplemental Table 4** for protein identities; upload table to <https://plothub.pages.dev/> for interactive visualisation.

In our study, we observed the detection of COWP8 in knockout TMT samples, where it was supposed to be absent. This unexpected detection can be attributed to reporter ion interference and co-isolation,[50] which are known limitations of TMT-based quantification in complex samples. Despite this challenge, we opted for TMT labelling due to its significant advantages for our specific research needs. Our in-house proteomic facility consistently achieved superior proteome coverage using TMT compared to label-free approaches. Furthermore, TMT offers a robust method for normalizing batch effects, which is particularly valuable given the labour-intensive nature of *Cryptosporidium* sample acquisition and preparation and the necessity of producing replicates in batches over extended periods. Ultimately, the enhanced proteome coverage, improved normalization capabilities, and increased experimental flexibility provided by TMT were deemed essential for the success this and future multi-batch proteomic study of *Cryptosporidium*, justifying our choice of this methodology.

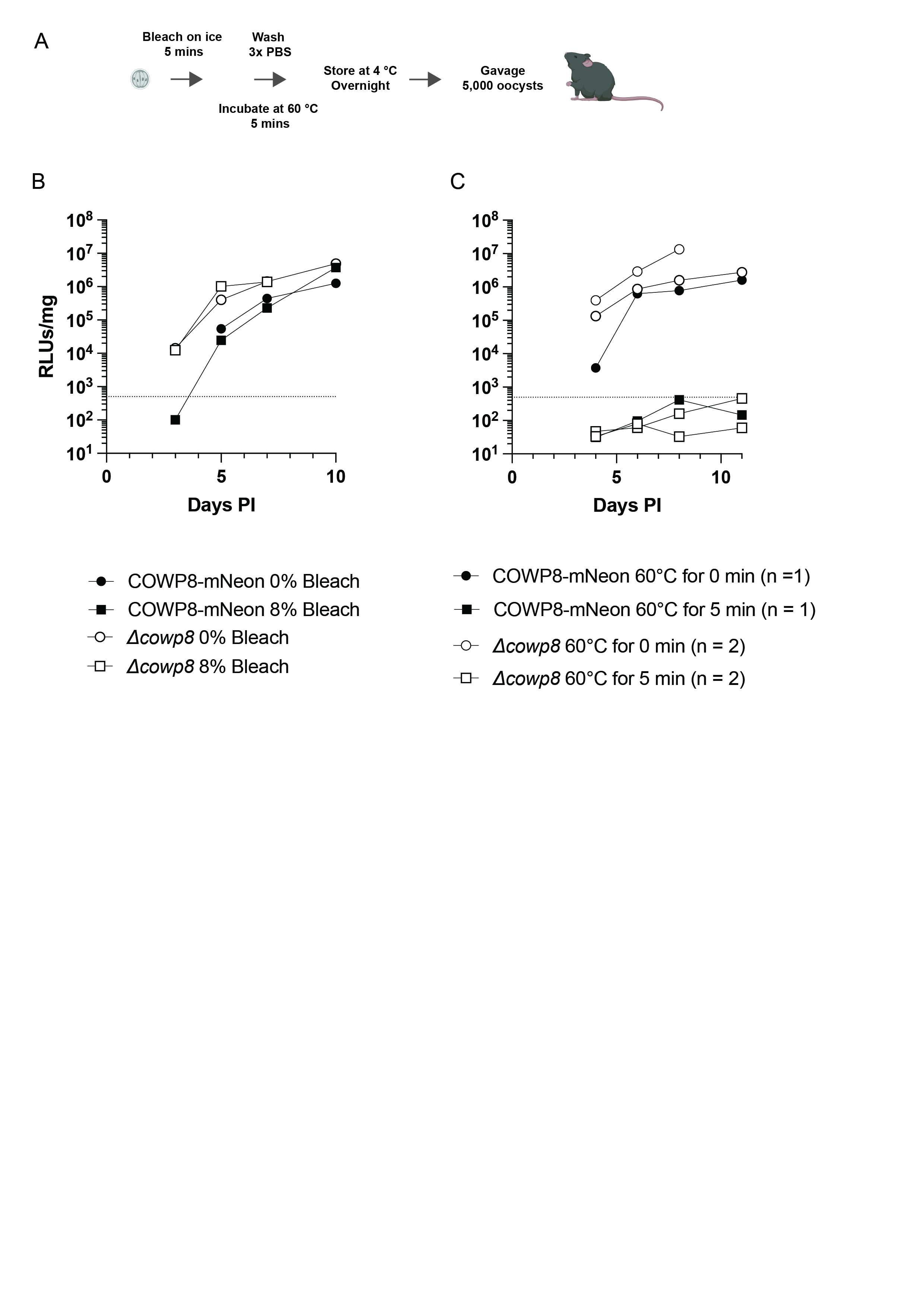

**Supplemental Figure 14. COWP8 Phenocopy COWP8-mNeon.**

**A**) Experimental design for testing oocyst resistance to bleach and temperature treatment. **B**) COWP8-mNeon (black) and *∆cowp8* strains (white) remain infectious after treatment with 8% bleach (squares). Average and SD of three technical replicates of one biological replicate. **C**) COWP8-mNeon (black) and *∆cowp8* strains (white,) are equally sensitive to heat inactivation (squares). Average and SD of three technical replicates of one biological replicate (representative experiment of 2 biological repeats performed for *∆cowp8*).

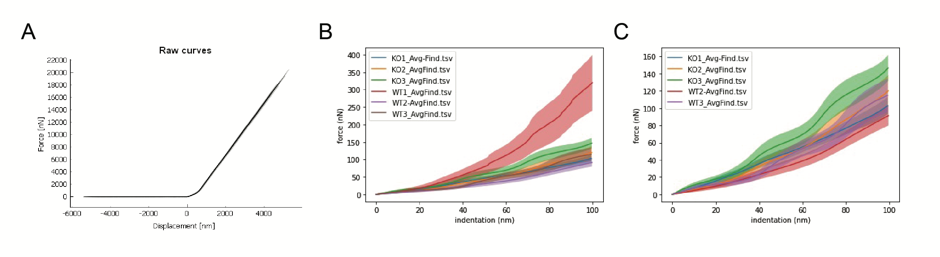

**Supplemental Figure 15. NanoIndentation of *Cryptosporidium* oocysts.**

**A**) Raw force measurements from NanoIndentation of wild type and *∆cowp8* oocysts. **B**) Force indentation plot containing all values (insert from **A)** in range of 0-100 nm of indentation. Three technical replicates performed each for wild type (red, purple, and lilac) and *∆cowp8* (blue, orange, and green). Included in the graph is initial replicate for wild type oocysts (“WT 1”). These values are determined to be outliers. This was the first sample generated during the first few measurements with the machine, when the techniques were being established. When the indentation graph is fitted to the Hertz Model to produce Young’s Modulus data, the outliers range is within that of polystyrene of the dish. Therefore, this sample represents the strength of the dish rather than wild type oocysts. Further optimisation was performed correcting the protocol. **C**) WT1 was therefore removed and revised plot of all replicates (excluding wild type biological replicate 1, WT1) are plotted.

**Supplemental Table 1.** Oligonucleotides used in this study.

| **COWP** | **Primer Name** | **Sequence 5´** ´ **-3´** ´ |
| --- | --- | --- |
| COWP2 | COWP2 gRNA F | GTTGGATTTGCTCAGGCAAAGTTT |
|  | COWP2 gRNA R | AAACAAACTTTGCCTGAGCAAATC |
|  | COWP2 Tag Homology F | GGTTAATTTTAAAAAAGAATCATTGCTTAAGATTTGCTCAGGCAAAGTTTAGATGTCCAATTGTAAGTGTGTCCAAGGGCGAGGAGGA |
|  | COWP2 Tag Homology R | CTTCAGTTTTCACTTGCTTAATACATTTTTCTCCTTCTTCCGTGAAACCCAATTAAGATAAAAAGAAAAACTTAATCGATACTATCCTACACGCC |
|  | 5´ In F COWP2 Tag | CCATTCCACGGTAAATTCCTGTCC |
|  | 5´ In R COWP2 Tag | CCATGCCCATCACGTCGGTGAACGCC |
|  | 3´ In F COWP2 Tag | GGCGGCTGTGCGAACGCATTCTGGC |
|  | 3´ In R COWP2 Tag | GCATGCACATGAAATATTTGTATGCGACC |
| COWP3 | COWP3 gRNA F | GTTGGCTAACTATAGAGAGTATTGA |
|  | COWP3 gRNA R | AAACTCAATACTCTCTATAGTTAGC |
|  | COWP3 Tag Homology F | CTAATTTCCAATACCAAAATCAAATGCAATATACAAGAAAAAATAAGTTCGTGTCCAAGGGCGAGGAGGA |
|  | COWP3 Tag Homology R | CGAACGGGCGCGCCTTTATGGCCGGTATAGCATGTATTTTTATAATGGCAAATTAAGATAAAAAGAAAAACTTAATCGATACTATCCTACACGCCACG |
|  | COWP3 at IMPDH Homology F | GCGCCAAATTCCTGTTATATGATAAATATTAATTAGATATCGACTATTTCGCATTTGGAATTGCATTGTG |
|  | IMPDH Homology R | GCTCATATGATTTAAATTTCAAAATGAAAAAACCAAAATAAATTTACAATTTAATATCACGGAAGGGGAATTAAGATAAAAAGAAAAAC |
|  | 5´ In F COWP3 Tag | GCTTAGTGGATTGGCGCCAACATATTATTTTATTCAAGTG |
|  | 5´ In R COWP3 Tag | CTACATTGAGTTGCTCCTGGTG |
|  | 3´ In F COWP3 Tag | GGTCAATGTGCTGCTCTTGAATTTGC |
|  | 3´ In R COWP3 Tag | CAAACTGGTGACTATATTTGTTTAACAAAC |
| COWP4 | COWP4 gRNA F | GTTGGCGTCGTATCTATATTCTGGT |
|  | COWP4 gRNA R | AAACACCAGAATATAGATACGACGC |
|  | COWP4 Tag Homology F | GTACTGGGACTTTTTGTAAAAGAACTGATTCGATCAAGCCCACACCTAGCTGTCCACCAGAATATAGATACGACG |
|  | COWP4 Tag Homology R | GCATACTTGGATTATGCTTGCGAGAGTCCAGGCTCTAATGAATGGTGAGGAATTAAGATAAAAAGAAAAACTTAATCGATACTATCCTACACGCCACGAAC |
|  | COWP4 at IMPDH Homology F | GCGCCAAATTCCTGTTATATGATAAATATTAATTAGATATCGACTATTTCAAAAATTTAGTTTCATTATC |
|  | 5´ In F COWP4 Tag | GCTTAGTGGATTGGCGCCAACATATTATTTTATTCAAGTG |
|  | 5´ In R COWP4 Tag | TGTTGTCCTCCTCGCCCTTGGACACTAAATTCAGATTTGTTCTAATGTTTG |
|  | 3´ In F COWP4 Tag | GCCGGTAGAGACTGGCTTCTTTTAGG |
|  | 3´ In R COWP4 Tag | CAAACTGGTGACTATATTTGTTTAACAAAC |
| COWP5 | COWP5 gRNA F | GTTGGTCACCTTCTTGGTCTAGGTG |
|  | COWP5 gRNA R | AAACCACCTAGACCAAGAAGGTGAC |
|  | COWP5 Tag Homology F | ATAGAAGAATAGGAGATCCCAGCTTAAATGTATATCCTCAAAACATTAAAACAGCACCTAGACCAAGAAGGGTGTCCAAGGGCGAGGAGGA |
|  | COWP5 Tag Homology R | AGATCCTGGTATATTCTTTAATCTCTGAGCAAAACTAATGGGCCTTATTGAATTAAGATAAAAAGAAAAACTTAATCGATACTATCCTACACGCCACG |
|  | COWP5 at IMPDH Homology F | GCGCCAAATTCCTGTTATATGATAAATATTAATTAGATATCGACTATTTCTACTATATACCGGTATTCATG |
|  | 5´ In F COWP5 Tag | GCTTAGTGGATTGGCGCCAACATATTATTTTATTCAAGTG |
|  | 5´ In R COWP5 Tag | GCTCTGGAGCTGATAACTTTACTCCAACGC |
|  | 3´ In F COWP5 Tag | GCCGGTAGAGACTGGCTTCTTTTAGG |
| COWP6 | COWP6 gRNA F | GTTGGCTTATGATCTAATTTAATAG |
|  | COWP6 gRNA R | AAACCTATTAAATTAGATCATAAGC |
|  | COWP6 Tag Homology F | AGCAAAATACACGAGAAGCACCATGGAAAAAATACTAGGACTGCAAATAGTCACTATATGGTATCAAAAGGAGAAGCGGTAATAAA |
|  | COWP6 Tag Homology R | ATTTAATACAAGTTTAGTTCGTTATTCGAAGTACTCAAAATCTTATGATCAATTAAGATAAAAAGAAAAACTTAATCGATACTATCCTACACGCCACGAACC |
|  | 5´ In F COWP6 Tag | GTCCCACCAAGCGGCCTATGTACAGC |
|  | 5´ In R COWP6 Tag | GTCTGCTAAGTATCTACCTCCGTCC |
|  | 3´ In F COWP3 Ta6 | GGCGGCTGTGCGAACGCATTCTGGC |
|  | 3´ In R COWP6 Tag | GAATGATCCAGTCTAAGTATCGAATACTC |
| COWP7 | COWP7 gRNA F | GTTGGAAAGACTTCTCATATAAAC |
|  | COWP7 gRNA R | AAACGTTTATATGAGAAGTCTTTC |
|  | COWP7 Tag Homology F | GCTTCCAACAGTATGCGCCTGCTCCAAGGTTTACAAATAGTCCATCTTACGCAGTTTATATGAGAAGTCTTTCGGAAGAAAAAAATGAGTTAATGGTATCAAAAGGAGAAGCGGTAATAA |
|  | COWP7 Tag Homology R | GAAGCTATATTTTATTGCTAAAAGATCACTCCACAATTAACATATCTAACAATTAAGATAAAAAGAAAAACTTAATCGATACTATCCTACACGCCACGAACCAATTGTCAGAAGAATTCG |
|  | 5´ In F COWP7 Tag | GGTGGAAGGTATCCTCCATATGAGAG |
|  | 5´ In R COWP7 Tag | GTCTGCTAAGTATCTACCTCCGTCC |
|  | 3´ In F COWP7 Tag | GGCGGCTGTGCGAACGCATTCTGGC |
|  | 3´ In R COWP7 Tag | GTAGCACAAAACATGTATGCATGCGG |
| COWP8 | COWP8 gRNA F | GTTGATTCAAACTACAATAACTGG |
|  | COWP8 gRNA R | AAACCCAGTTATTGTAGTTTGAAT |
|  | COWP8 Tag Homology F | GTTGTTGTACAACCAACAATGGCTTCACAACAACAAGAAATTATTCAAACTACAATAACTGGTGcACGTCATCATCATCATGTGTCCAAGGGCGAGGAGGA |
|  | COWP8 Homology R | GAAGAAGAGTGTAGTAAATGGATGAATTATCTTAATTCGGTAATCCCTCCAATTAAGATAAAAAGAAAAACTTAATCGATACTATCCTACACGCCACGAAC |
|  | 5´ In F COWP8 Tag | GATGTGCACGTTACATGCATGCATTC |
|  | 5´ In R COWP8 Tag | CGTCAGCGAGTTGGTCATCA |
|  | 3´ In F COWP8 | GGCGGCTGTGCGAACGCATTCTGGC |
|  | 3´ In R COWP8 | GAAATGGAGGAGCTTCTAAAGCATGATG |
|  | COWP8 KO Homology F | GAAATTAGTCATAAGAGGAAGTACACCAGATACATGGTAGTTATAAAACATGGGGAAACTAAATATACTGAAATTCGGTAG |
|  | COWP8 ORF F | CTGACGGTCAAATGTGCAGTGGCTC |
|  | COWP8 ORF R | GGACATCCTAATTCACCAGGAACAG |
|  | 5´ In F COWP8 KO | GATGTGCACGTTACATGCATGCATTC |
|  | 5´ In R COWP8 KO | GTGCCATAGTGCAGGATCACCTTAAAG |
| COWP9 | COWP9 gRNA F | GTTGAGATAAGAATAACCCTCAAA |
|  | COWP9 gRNA R | AAACTTTGAGGGTTATTCTTATCT |
|  | COWP9 Tag Homology F | GGAAGAACTGTGGTTCCTTCTTATTC |
|  | COWP9 Tag Homology R | CCACATGAATGCATGCATGTAACGTGCACATCAAAATTTGCATACCGGTAAATTAAGATAAAAAGAAAAACTTAATCGATACTATCCTACACGCCACGAACC |
|  | 5´ In F COWP9 Tag | CATGAATCTACCGAACATAATGTGAA |
|  | 5´ In R COWP9 Tag | CCATGCCCATCACGTCGGTGAACGCC |
|  | 3´ In F COWP9 Ta6 | GGCGGCTGTGCGAACGCATTCTGGC |
|  | 3´ In R COWP9 Tag | CGGAGACTGTTGAAACTGCTTGTTGTTG |
| Controls | TK ORF F | ATGGCAAAATTATACTTTTACTATTCAGCAATGAATGC |
|  | TK ORF R | TTAGAAATTGTATTCTTCACAATTAATTATATGATGTTTTCTGC |
|  | a-tub F | CTAGTTATCCTTGTTCATTGAATTCTC |
|  | a-tub R | TGAGCTCAAAAATATAAGATGGCAC |
|  | IMPDH ORF F | GATGTAATTGTTGGGAATGTTGTAACAGAAGAAGCAAC |
|  | IMPDH ORF R | GCAGGATCTTAGTCCTCCAACAAGCTGATATACTACACCTTCCATTTCACC |

**Supplemental Table 2.** Plasmids created for this study. Links to full sequence available through link.

| **Purpose** | **Plasmid name** |
| --- | --- |
| mScarlet-I tagging | [pLIC-mScarlet-I-3Xc-myc_aldo 3´ utr_ENNE](https://benchling.com/s/seq-nGebYhPHY3pwwQE8MxLt?m=slm-HYkPHRjJH5m8j0Z3XuAJ) |
| mScarlet-I knockout | [EnoP_mScarlet-I-ENNE](https://benchling.com/s/seq-KCnDG09dRQe7RqfvUSJR?m=slm-DBxGDhCzrd86KNSbPMUK) |
| COWP4-C-term tagging | [*Cp*COWP4 C-term-mNeon-3XHA_ENNE](https://benchling.com/s/seq-7yo2dj2E2KIMG81mIXxp?m=slm-ABwgOjSrps6yM3lGu8Is) |
| COWP9-C-term tagging | [*Cp*COWP9 C-term-mNeon-3XHA_ENNE](https://benchling.com/s/seq-gQU4QAKhTls1uGoBRI45?m=slm-5UDJcbF6fU39RhHxedEt) |
| COWP3 expression | [Cowp3P_COWP3ORF-mNeon-3XHA_ENNE](https://benchling.com/s/seq-8DyxXhWXcfDibsb1kFNY?m=slm-OzrqYN73iMLc0TmPJk5W) |
| COWP4 expression | [Cowp4P_COWP4ORF-mNeon-3XHA_ENNE](https://benchling.com/s/seq-3oLCypzWRkoLmiJ58J8x?m=slm-00gm58LW2DUyPL5QEgo2) |
| COWP5 expression | [Cowp5P_COWP5ORF-mNeon-3XHA_ENNE](https://benchling.com/s/seq-mnZ1gjD48dSdy5mnKE4W?m=slm-krZLoM5MVdrAKIl7RzNW) |

**Supplementary Table 3.** Antibodies and dyes used in this study.

| **Antibody name** | | **Supplier**  (Catalog #) | **Purpose** | **Working dilution** |
| --- | --- | --- | --- | --- |
| **Anti-HA (rat)** | | Sigma Aldrich (11867423001) | ImmunoEM | 1:300 |
| **Anti-c-myc (mouse)** | | ProteinTech  (10828-1-AP) | ImmunoEM | 1:100 |
| **1A5 (mouse)** | | Sibley Lab | IFA | 1:500 |
| **1B5 (mouse)** | | Sibley Lab | IFA | 1:500 |
| **DAPI** | | Sigma Aldrich (10236276001) | IFA |  |
| **Phalloidin-647** | | Abcam (ab176759) | IFA | 1:1000 |
| **Hoechst 33342** | ThermoSci (62249) | | IFA | 1:2000 |

**Supplementary Table 4. Differential Expression Analysis of *C. parvum* Genes.**

The table (attached as csv file) presents a summary of the differential expression analysis results. Each row corresponds to a unique gene characterized by the following attributes: “Gene_acc”: a unique integer identifier. “Gene_id”: The gene identifier from CryptoDB. “logFC”: The logarithm (base 2) of the fold change in TMT intensity between the WT (Wild Type) and KO (COWP8 knockout) samples as computed by the limma R package. “log_AveExpr”: The logarithm (base 2) of the average gene expression level across all samples as computed by the limma R package. “FDR”: The False Discovery Rate-adjusted p-value. Desc: The gene description from CryptoDB. “WT_1” to “WT_4”: The MaxQuant quantified TMT intensity levels for each of wild-type replicates, normalized and fed to the limma package. “KO_1” to “KO_4”: TheMaxQuant quantified TMT intensity levels for each of the overexpression condition, normalized and fed to the limma R package. For enhanced visualization and exploratory data analysis, the table has been formatted to be compatible with the interactive plotting tool available at https://plothub.pages.dev/. This online resource provides an intuitive graphical representation of the data, allowing for immediate visual assessment and interpretation.
